# Core variability in substitution rates and the basal sequence characteristics of the human genome

**DOI:** 10.1101/024257

**Authors:** Aleksandr B. Sahakyan, Shankar Balasubramanian

## Abstract

Accurate knowledge on the core components of substitution rates is of vital importance to understand genome evolution and dynamics. By performing a single-genome and direct analysis of 39894 retrotransposon remnants, we reveal germline sequence-dependent nucleotide substitution rates that can be assigned to each position in the human genome. Benefiting from the data made available in such detail, we show that a simulated genome, generated by equilibrating a random DNA sequence solely using our rate constants, exhibits nucleotide organisation close to that in the human genome. We next generate the germline basal substitution propensity (BSP) profile of the human genome and show a decreased tendency of moieties with low BSP to undergo somatic mutations in many cancer types.

## Introduction

The stability, organisation and dynamics of genomes are key factors that influence the molecular evolution of life [1]. Single-nucleotide mutations and their subsequent fixation in a population occur an order of magnitude more frequently than common insertions/deletions [2, 3], hence are major contributors in defining the possibilities (sampling pathways) for a genome to evolve. A thorough understanding of the descriptors that govern single-nucleotide substitutions is thus essential to comprehend genome dynamics and its connection to the underlying single-nucleotide processes.

For a given genomic position and *i→j* nucleotide substitution, the substitution rate, as expressed by the rate constant *r*_i,j_, can be presented as a single-base average value 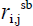 and fluctuations contributed by short-range context (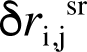), CpG-associated (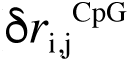), long-range (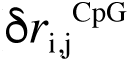), gene/functional (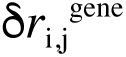), and specific (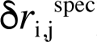) effects:

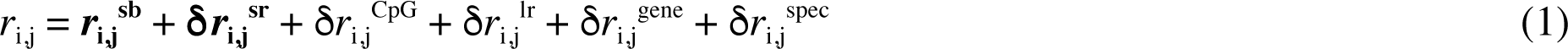

The 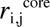 term can be estimated through genomic averages for the individual *i→j* substitution rates, and has been reported for the genomes of human [4, 5] and other species [6-8]. By investigating the aggregation patterns in substitution frequencies, it was shown that the *r*_i,j_ variation is subjected to two distinct short-range (<10 nt) and long-range (>1000 nt) effects [9, 10]. In the equation above, the short-range effect is captured through the 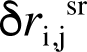 term and describes the totality of the intrinsic properties and sequence-dependent interactions of DNA with overall mutagenic and reparation processes in a given organism [11]. The better-studied substitution patterns at a CpG context [12-14] are separated in the 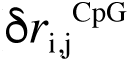 term, since besides having a specific short-range dyad context, CpG mutations also depend on a number of regional factors that alter the epigenetic targeting of the CpG sites [10, 15-18]. Many relatively recent studies have shed light on the 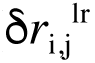 variation caused by the regional effects that depend on a long-range (megabase) sequence context through mechanisms, such as recombination and GC-biased gene conversion [1, 4, 19, 20], transcription-coupled biased genome repair [21] and instability [22], chromatin organisation [23], replication-associated mutational bias [24] and inhomogenous repair [25], differential DNA mismatch repair [26], non-allelic gene conversion [27], and male mutation bias [28]. The term 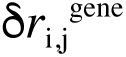 captures the change in substitution rates in genes and other functional elements under strong selection bias and reflects observations such as the increased neutral substitution rates in exons [29, 30] and the possible reduction of the mutation rates in X-chromosome [31]. 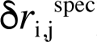 holds the highly specific increase or decrease in substitution rates governed by targeted hyper- and hypomutations [32] present, for example, in the genes of immune system.

Herein, we obtain the core components (**equation 2**) of the spontaneous single-nucleotide substitution rates via the direct analysis of 39894 L1 mobile DNA remnants [33] in the same, human, genome (a single-genome approach).

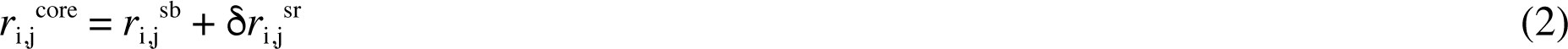

Our **tr**ansposon **e**xposed **k** (Trek) method provides the 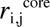 rates at single-nucleotide resolution in L1, where we demonstrate sufficient sequence variability to cover a wide-range of sequence contexts. We use this coverage to determine the core rate constants for all possible nucleotide substitutions (3 per position) at every single 3.2 billion position in the human genome. The Trek method reveals the 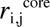 variation in a model-free manner and at a level beyond accounting for only the two immediate neighbouring nucleotides [34]. Furthermore, we make our dataset, holding the time-dependent rate constants for individual substitutions that account for up to 7-mer sequence context effect, publically available. Importantly, we demonstrate that 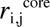 values alone can generate a sequence, from first principles, starting from a random DNA sequence, whose key features reflect the oligomeric organisation of the actual human genome. Next, we calculate the basal substitution propensity profile of the human genome, evaluating the core predisposition to single-nucleotide substitutions. We outline the decreased frequency of the sequence motifs that are stable in germline among the sites linked to somatic cancer mutations.

## Results and Discussion

### Revealing the Core Single-Nucleotide Substitution Rates

The repetitive occurrence of mobile DNA elements in different regions within the same genome [33] provides the opportunity to obtain the core 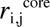 substitution rate constants that account for the 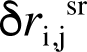 immediate effects of neighbouring nucleotides. After the initial inactivation at different time epochs [35-38], individual remnants of many transposon subfamilies within a genome have been subjected to largely the same overall mutagenic and proofreading conditions as the rest of the genome [39], hence can also serve as markers of 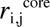 substitution rates applicable to genomic sites that share the immediate sequence-context. For the purpose of this study, we have used the hominoid lineage of the L1 (LINE-1) retrotransposons, spanning 3.1 to 20.4 myr (million years) of age [36]. The constituent subfamilies of the lineage are L1PA5, L1PA4, L1PA3, L1PA2 and the most recent L1Hs. Their respective age and the number of insertions in the human genome are presented in **supplementary table S1**. The choice was made through the following reasoning. The L1 elements have a long (∼6k nt) sequence without extended repeats like in the LTR elements [33]. This enables their robust mapping on a chosen template and provides essential local sequence variability around different nucleotide positions within L1 elements. There are distinct L1 subfamilies that were active at different time epochs, with detailed molecular clock analyses available [35-38] to reveal and, importantly, validate the age of each subfamily. They are well-represented and, unlike other classes of transposable elements, are uniformly scattered across mostly the intergenic regions of the human genome [33, 40, 41]. Unlike SINEs and LTRs, LINE sites show very low level of RNA polymerase enrichment, as a marker of transcriptional association, in normal tissues [42]. The selected most-recent subfamilies are sufficiently young [36] a) to enable an unambiguous identification of the genomic coordinates of the borders for the remnants; b) to assume that each position in those elements would be unlikely to undergo repeated substitutions over the studied period of their existence as remnants in the human genome (see **Methods**); c) to attribute a time-invariance to the rates during the analysed period of substitution accumulation [4, 43, 44]. The young L1 subfamilies have most of their remnants coming from the genomic regions with G+C content close to the genomic average value [40] (see **supplementary fig. S1**). Finally, many matching positions in our studied five L1 representatives share the same consensus bases, hence, such positions are not polymorphic due to adaptive pressure and can serve as internal references for inferring the 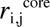 rates.

The Trek methodology of obtaining 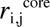 rates, along with the considerations for filtering out the possible selection and non-neutral substitution sites, is presented in **fig. 1** with further details in **Methods** and **supplementary fig. S2**. The acquired data on the full set of position-specific substitution rates are presented in **fig. 2**, to highlight the revealed variation per substitution type.

**Fig. 1.**
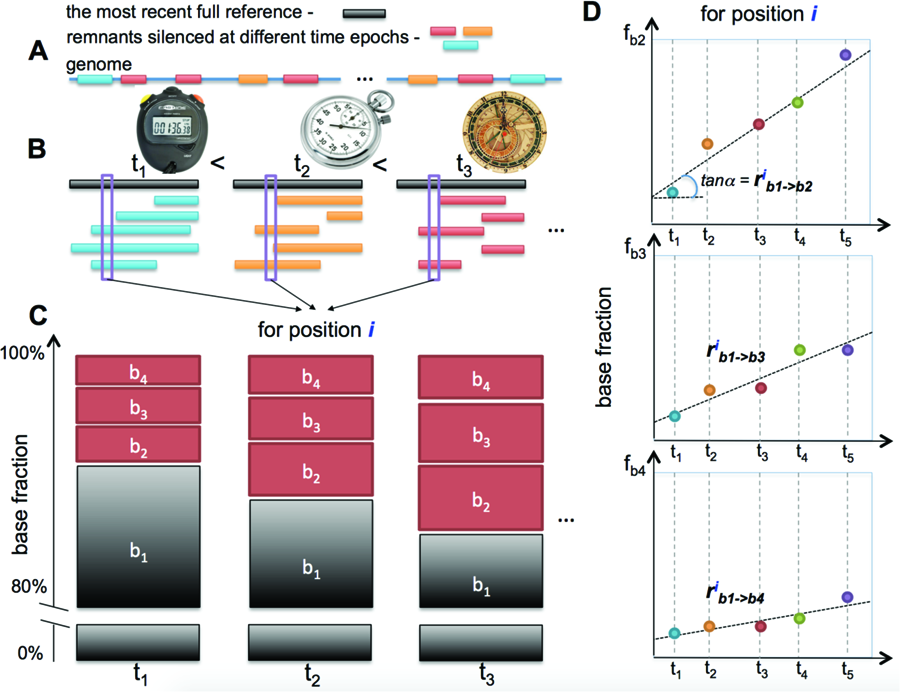
Trek methodology of determining the core single-nucleotide substitution rate constants. (**A**) The Trek approach is applicable to a genome containing multiple remnants of retrotransposon subfamilies silenced at different time epochs. We can consider those subfamilies as substitution counters that had different resetting ages (**B**). The full consensus sequence of the most recent subfamily is taken as a reference (**A**). The remnants are then grouped by their age and fully mapped onto the reference sequence (**B**). For each position *i* in the reference sequence, the fractions of the four bases in all the time groups are calculated (**C**). The comparison of these fractions coming from individual base types across different time periods enables a linear model fitting, through which we can reveal the rates for the substitutions into the *b*_2_, *b*_3_ and *b*_4_ bases from the consensus ( *b*_1_) state of the given position (**D**). The steps **C** and **D** are repeated for all the positions in the reference sequence, producing single-nucleotide resolution core substitution rate constants with a sequence-context dependency as sampled in the reference sequence of the mobile element. To assure the high quality and neutrality of the retrieved rates, we accounted for the sites in the reference sequence that had at least 700 mapped occurrences in each time group (**B**), with the same wild-type variant being always the prevalent one (more than 80%) in each subfamily (**C**) and producing a Pearson's correlation coefficient of at least 0.7 in the time-evolution plots (**D**).

**Fig. 2.**
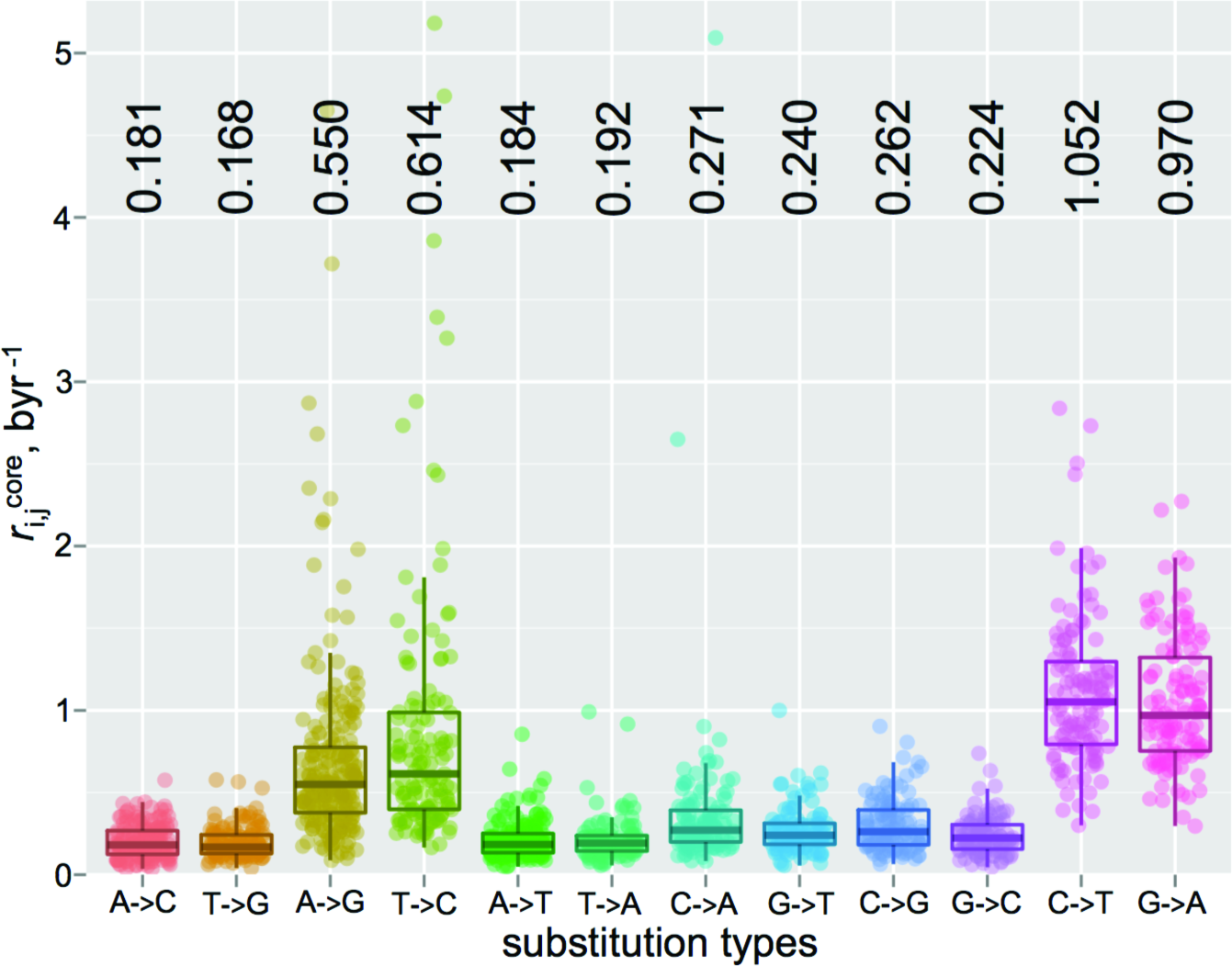
Transposon exposed (Trek) 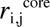 substitution rate constants of the human genome. The boxplots are shown for each *i→j* substitution type inferred from the hominoid L1 elements spread across the human genome. Each point comes from a specific position in the L1 element, reflecting the substitution rate constant averaged across multiple occurrences of that specific position with the same sequence-context in multiple regions of the genome. The complementary *i→j* pairs are plotted in adjacency. The median values of the overall substitution rates in byr^−1^ (billion years) unit, averaged across the varying sequence-context within the L1 elements, are shown on the top.

A total of 661 positions, at the 3’ side of the L1 elements, passed our robustness checks (see **Methods**) and were thus employed to infer the corresponding 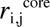 values from the analysis of all the young L1 remnants in the human genome. We recorded the data in the Trek database that contains a set of well-defined 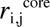 constants (see below for the extent of sequence context coverage in the Trek database) capturing the influence of the unique arrangement of neighbouring nucleotides at those positions. Owing to the nature of the selected L1 elements, as discussed above, and the Trek procedure design (**fig. 1**), we expect the absence of the 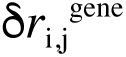 contribution, the elimination of 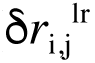 at the averaging stage (**fig. 1B**) and the removal of the 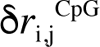 and 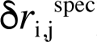 effects through our robustness checks embedded within the Trek procedure (see **fig. 1C**,**D** and **Methods**). Therefore, our method provides the 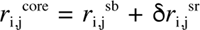 core variation (**fig. 2**) of the substitution rates at around the 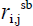 genomic average values for each *i→j* base substitution. If the above is correct and our method indeed results in 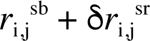 values, further averaging of the core 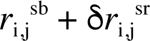 rates (median values shown in **fig. 2**) should give us the single-base 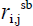 genomic average substitution rates, cancelling out the remaining 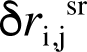 contribution. In fact, the comparisons of our Trek-derived 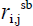 with two published datasets that report on the genomic average 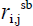 rates [4, 5] show an excellent correlation (**supplementary fig. S3**, Pearson’s R>0.99) confirming the absence of any bias and unusual substitution rates in the time-accumulated substitutions at the L1 sites that pass the Trek procedure. The genome simulation, described later in this work, provides an additional validation for our rate constants. The 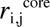 values (**equation 1** and **2**) for all possible *i→j* substitutions inferred for each of the eligible individual L1 positions are thus assumed to be common for any other sites in the genome that share the short-range sequence context.

### The Influence Range of Neighbour Nucleotides

To apply the 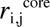 constants to the human genome, we first established the optimal length of a DNA sequence (k-mer, where k is the length of the sequence) capturing most of the influences that modulate substitution rates of the base at the centre. For this, we evaluated the power of the knowledge of the neighbouring arrangement of nucleotides in predicting the 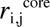 constants for each of the twelve *i→j* substitution types, where *i* and *j* are the four DNA bases. We built test predictors for individual substitution types via a tree-based machine learning technique, while using varying lengths of sequences centred at the positions where the rate constants were to be predicted (see **Methods**). The aim of the machine learning procedure was to establish the optimal sequence length to minimise the error in the predicted rate constants (**supplementary fig. S4** and **S5**). In agreement with prior evidence [9, 10, 45, 46], but now obtained for each individual *i→j* substitution type from Trek data, the optimal window was found to be 5-7-nt (both 5- and 7-nt resulting in comparable results for many substitution types) and was subsequently used as guidance for the direct mapping of the Trek rate constants from the L1 sequence onto any given human nuclear DNA sequence for the 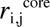 assignment.

### Mapping the Trek 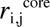 Data on any DNA Sequence

The upper 7-nt size window for determining the single-nucleotide substitution rate constants at the central base accounts for 3 upstream and 3 downstream bases relative to each nucleotide position. Our substitution positions that pass the Trek criteria capture 636 unique 7-mers out of the possible 16384 (4^7^). Therefore, for many loci in the human genome we need to use a smaller window (< 7-mer) as a match criterion to assign to one of the Trek rate constant sets. By trimming the size of the k-mer to 5, hence accounting for 2 upstream and 2 downstream bases, we cover 404 unique sequences out of possible 1024 (4^5^). Further reduction of the size to 3, allows having data for 56 unique triads out of 64 (leaving out only the CpG containing triads, see below). For the single-base case (1-mers), where we average out all short-range neighbour effects and longer-range sequence variability, we obtain data for all the 4 bases and 4×3 possible substitutions as shown in **fig. 2** (the median values on the top of the figure). The coverage of the longer k-mers is however increased nearly twice when we account for the strand-symmetry, as described in **Methods**. Please note, that for each unique k-mer we obtained 3 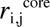 constants via the described analysis of a large pool of L1 remnants from different genomic loci.

With the above considerations, we created a program (Trek mapper) to produce the full range of 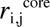 core substitution rate constants for any sequence, accounting for the context information within up to the 7-mer window and pulling the matching core data from the Trek database. Should a representative match be absent with the full 7-nt long sequence, the window around the given position in a query sequence is shortened into the longest variant possible (out of the 5-nt, 3-nt or 1-nt lengths) with a full match in the Trek database (see **Methods** and **supplementary fig. S6**). In this way, for all the possible 16384 7-mers, our Trek database reports 49152 rate constants (3×16384), of which 3168 (6.4%) account for the 7-mer context, 23232 (47.3%) account for the nested 5-mer context, 17120 (34.8%) for 3-mer and only 5632 (11.5%, CpG containing sequences) constants do not account for any context effect on the central base, since we eliminate those by design, due to the 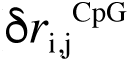 contributions. We thus produce an unprecedented dataset that reports, and makes publically available (**supplementary data 2** and **data 3**), the time-dependent 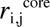 rates for all individual *i→j* substitutions accounting for the context effects beyond the 64 triads [34]. If we consider only the unique values in the Trek database, we report 2078 unique rate constants (taking into account different extent of averaging, where multiple entries are present for the different context ranges), of which 1208 (58.1%), 782 (37.6%), 85 (4.1%) and 3 (0.1%) entries account for 7-, 5-, 3- and 1-mer contexts respectively. The 1-mer averaged data were used for the k-mers in that contain either C or G bases of a CpG dyad at the centre, to assign the overall substitution rate constants by the Trek mapper. This was done since none of the CpG sites in the L1 elements passed our robustness checks, due to the targeted epigenetic control (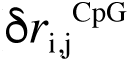) of the single-nucleotide substitutions there [12-14], which were also non-uniform with time (active targeting, 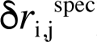, while in the viable epoch for each L1 subfamily) and were present to silence the active retrotransposons.

Our current data are for the human nuclear genome. However, the general approach for obtaining 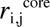 constants is applicable to any organism where the genome contains a set of well-characterised and related young mobile elements silenced at different time epochs and without notable context bias.

### 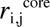 Rates and the Oligomeric Landscape in the Human Genome

The full set of sequence-dependent human 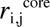 substitution rates (all 3 constants per position) enabled us to perform a sophisticated *in silico* evolution of a random DNA sequence, guided solely by our 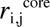 values. We started from a random sequence of 5 million (mln) nt with a G+C content of 60% (substantially greater than the 40.45% G+C content for the human genome). We performed random nucleotide substitutions weighted by Trek-inferred probabilities (see **Methods, supplementary fig. S7** and **supplementary video 1**), where, after each cycle, the substitution rate constants were updated for the sequence positions that were either mutated or fell within the influence zone of the performed substitutions. The simulation was continued until the overall G+C content of the simulated sequence became constant (see **fig. 3A-C**).

**Fig. 3.**
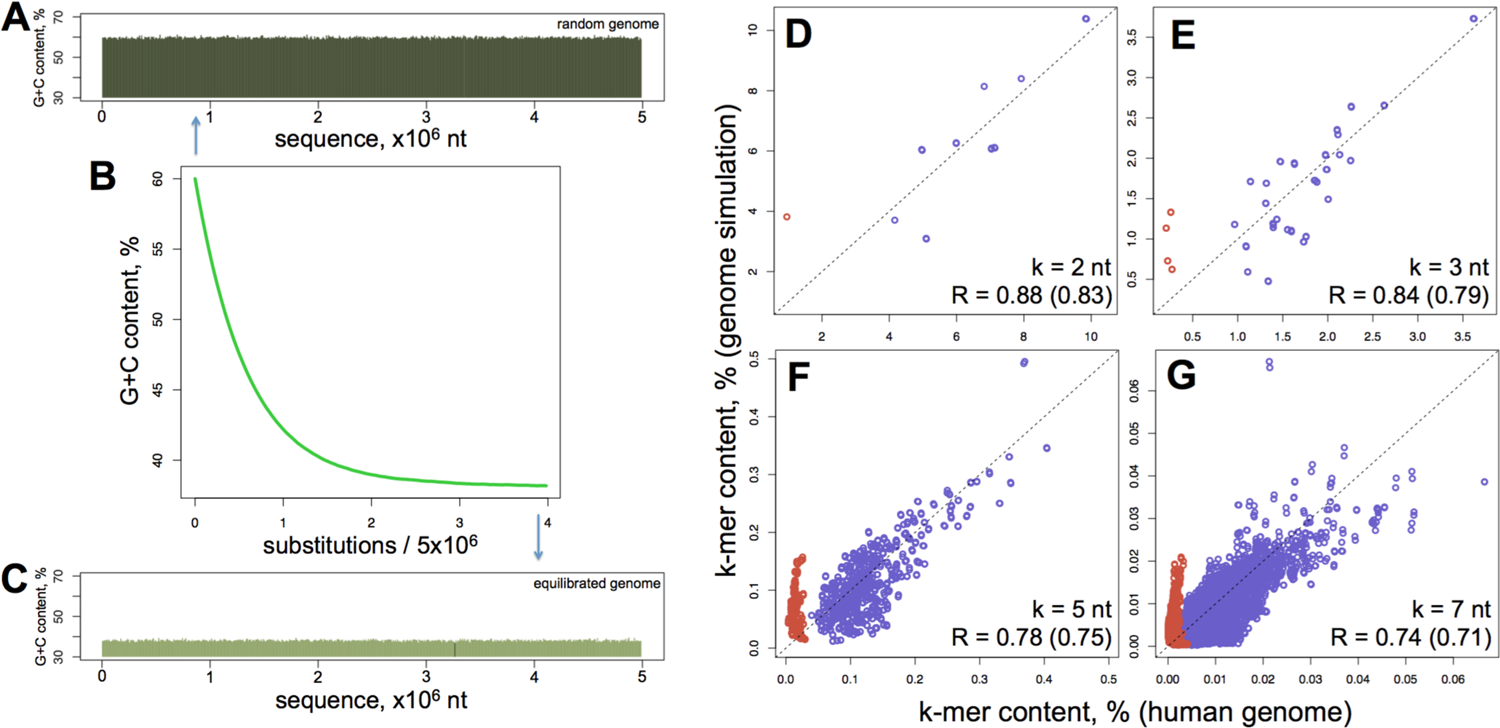
Comparison of the *in silico* evolved and actual human genomes. (**A**) The 5-mln-nt starting sequence is randomly generated with 60% G+C content. The sequence is then neutrally evolved using 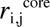 values only, until the base-compositional equilibrium is established (**A-C**). This was reached after about 20 mln substitutions (or an average of 4 substitutions per site (**B**). The equilibration converges faster when we start from a sequence with lower G+C content. The plots **D-G** show the correlation of the k-mer contents in the equilibrated genome with the corresponding content in the real human genome. The lengths of the k-mers along with the correlation coefficients are shown on the bottom right corners of the plots. Two correlation coefficients are shown with the exclusion and the inclusion (the value in the bracket) of CpG containing oligomers (red points in the plots). The dashed lines depict the diagonals for the ideal match of the k-mer contents.

The simulation converged to generate a sequence with the A, T, G and C compositions of 30.91, 30.90, 19.06 and 19.13% respectively. Note, that these values are much closer to the A, T, G and C compositions of the repeat-masked human genome of 29.75, 29.79, 20.24 and 20.22% respectively (see **Methods**), being just slightly AT rich. Furthermore, the simulated genome captures the contents of different individual oligomers (k-mers) in the human genome. The data for all the possible 16 dyads, 64 triads, 1024 pentads and 16384 heptads are presented in **fig. 3D-G** and show a significant (see the correlation coefficients) correlation between the compositional landscapes of the Trek-simulated genome and the actual human genome. Regardless of the starting composition of the initial DNA sequences, our simulations always equilibrated to a state with similar oligomer (up to 7-mer) content. The k-mer contents shown in **fig. 3D-G** for the actual human genome were calculated from the repeat-masked version of the RefSeq human genome, where all the identified repeat elements, including the L1, were disregarded. This assured the removal of a potential bias due to the presence of L1 elements in the human genome. As 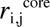 constants are free of the 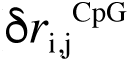 contribution (see above), the simulated genome produced higher alterations in representing the k-mer contents that have CpGs (red points in **fig. 3D-G**). These alterations directly demonstrate the contribution of 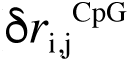 to the background compositional landscape of the human genome.

The correlations in **fig. 3** are from simulations where the rate constants were symmetrised according to the inherit strand-symmetry in double-helical DNA (see **Methods**). The results without such equalisation are still significant, though producing slightly worse correlation coefficients (**supplementary fig. S8**).

To confirm that the observed correlations for different k-mer contents (**fig. 3D-G**) present an improvement due to our sequence-context-dependent core substitution rates, rather than being a side effect, by a pure chance, in a sequence where the simulation makes only the single-base composition converge to that of the real human genome (such as in sequence generated using an ideal 4×4 single-nucleotide substitution rate matrix), we calculated the expected distribution of different k-mers in a genome with fully random base arrangement but with the exact human A, T, G and C overall base composition. In the complete absence of any sequence-context effects, the probability of the occurrence (fraction) of any k-mer in a sufficiently long sequence is equal to the product of the occurrence probabilities of their constituent bases. For instance, the probability of observing the AGT triad is the p_AGT_ = p_A_p_G_p_T_ product, where the individual p_i_ probabilities are the base contents expressed in fractions. The comparison of the k-mer fractions obtained in this way with the human genome data (**supplementary fig. S9**) shows a substantially reduced correlation (for the genomic 7-mer content, Pearson’s R=0.59 compared to 0.74 using Trek rates), supporting the attribution of the important role to the 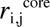 variability in shaping the compositional landscape of our genome.

### Basal Substitution Propensity Profile of the Human Genome

Our Trek mapper provides the full set of 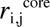 constants for each position in the whole human genome. Such data enables us to calculate the germline context-dependent basal substitution propensity (BSP) by taking the sum of the individual rate constants for the 3 possible substitutions at each base position, thus producing the core 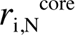 constant for the substitution of a given base *i* by any other base *N*. **Supplementary fig. S10** shows the BSP profiles calculated for the individual chromosomes (red) as compared with the whole genome profile (green), where most of the chromosomes exhibit the same overall distribution as the whole genome. Further grouping and analysis [47] of the unique sequences found in regions of different BSP for the whole human genome reveals motifs that tune the stabilities of the bases at the centre (see **fig. 4** and the caption for further elaborations). Note that the observed sequence-determined substitution biases can potentially contribute to the initial nucleation of a more extensive base-content pattern (isochore) formation in chromosomes [48, 49], which are prevalently driven by longer-range regional effects and caused variability in substitution rates.

**Fig. 4.**
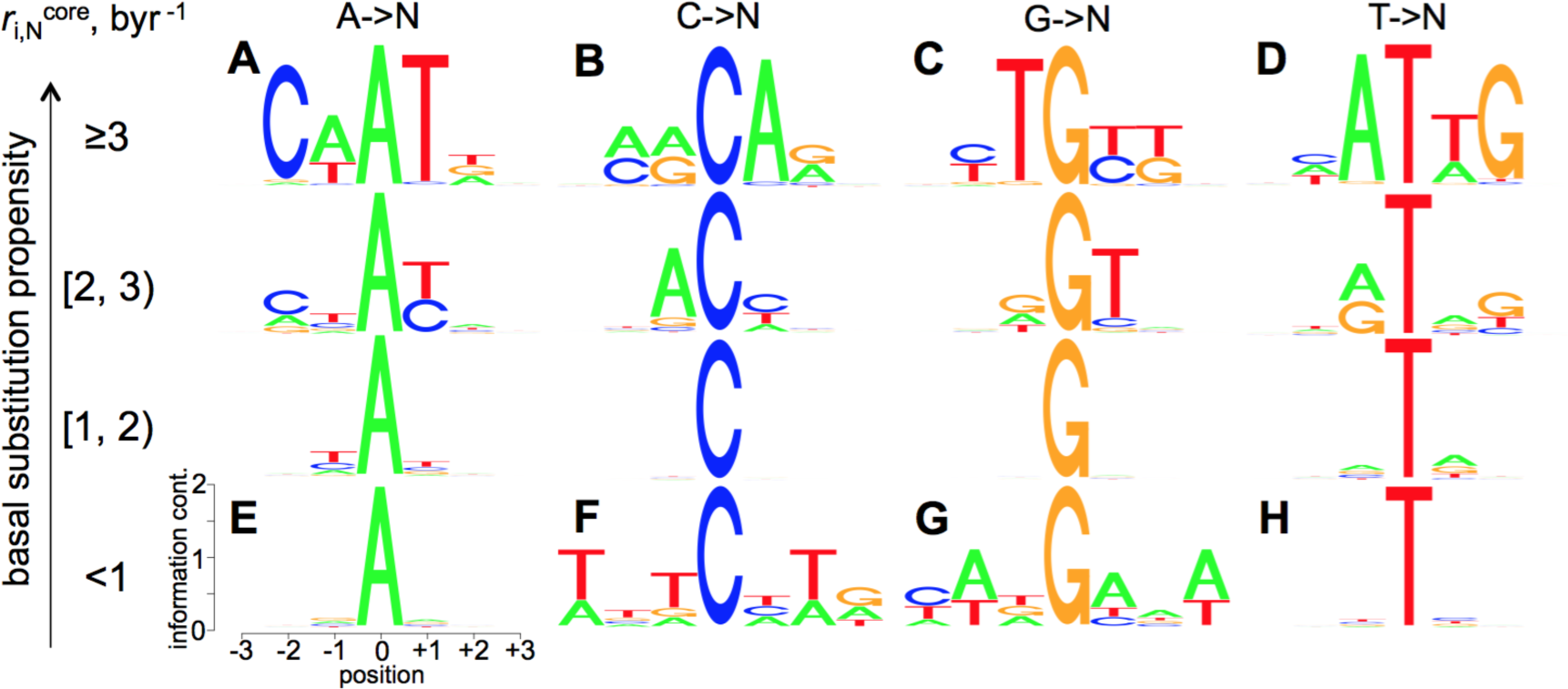
Sequence-context dependence of the 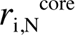 basal substitution propensity (BSP) constants. Sequence logos [47] are shown for all the unique 7-mer sequences grouped by the central base type (columns) and the category of the BSP range the sequences fall in (rows, BSP range is shown in byr^−1^ rate constants). The y-axes in the individual sequence logos show the information content in bits. The x-axes outline the neighbouring base positions relative to the central base. For each sequence, the BSP of the central base *(i)* depicts the sum of the core rate constants for the substitutions to the three other (non-*i*) bases, 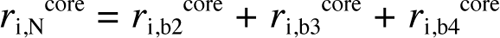. As can be seen from the plots, the bases A and T are highly mutable when the neighbouring positions are enriched in the same, A and T, bases (compare the logos **A** and **D** with **E** and **H**). The adjacent enrichment in A increases the BSP of C (**B**), and decreases the BSP of G (**G**) bases. Conversely, the adjacent enrichment in T increases the BSP of G (**C**) and decreases that of C (**F**) bases. Note, that our data are for 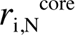 and do not include the methylation-driven increased mutation rates in CpG dyads [12-14].

### Basal Substitution Propensity Profile and Cancer-Linked Somatic Mutations

A recent study [50] suggested that cancer, with its multi-etiologic nature, is primarily linked to random mutation events upon cell division/DNA replication. To this end, genomic sites with higher intrinsic BSP (lower stability) might exhibit a higher prevalence of cancer-related genome alterations, as compared to sites of lower intrinsic BSP (higher stability), should the germline and cancer-linked somatic mutations share common mechanisms. Although the sequence context signatures of cancer mutations and their variation across different cancer types is out of the scope of the present work and is covered in detail elsewhere [51-54], here we examined the simple relationship between our calculated germline BSP values and the observed cancer-associated somatic mutations accessed via the annotated COSMIC database of somatic mutations in cancer [55] (see **Methods**). Since the Trek data are for the core spontaneous substitutions, we restricted the analysis to the non-coding and non-polymorphic (not identified as SNP) point mutations (6 mln) in cancer. By mapping these sites to the human genome and retrieving the sequence-context information (7-nt long sequences centred at the mutation points), we processed the data with Trek mapper and obtained the BSP 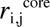 profile for the non-coding sites detected in human cancer. The outcome in **fig. 5**, overlapped with the whole-genome BSP profile, shows that stable sites in the human genome, assigned by the Trek mapper to have 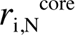 below 1.13 byr^−1^, are significantly less likely to undergo somatic mutations in cancer.

**Fig. 5.**
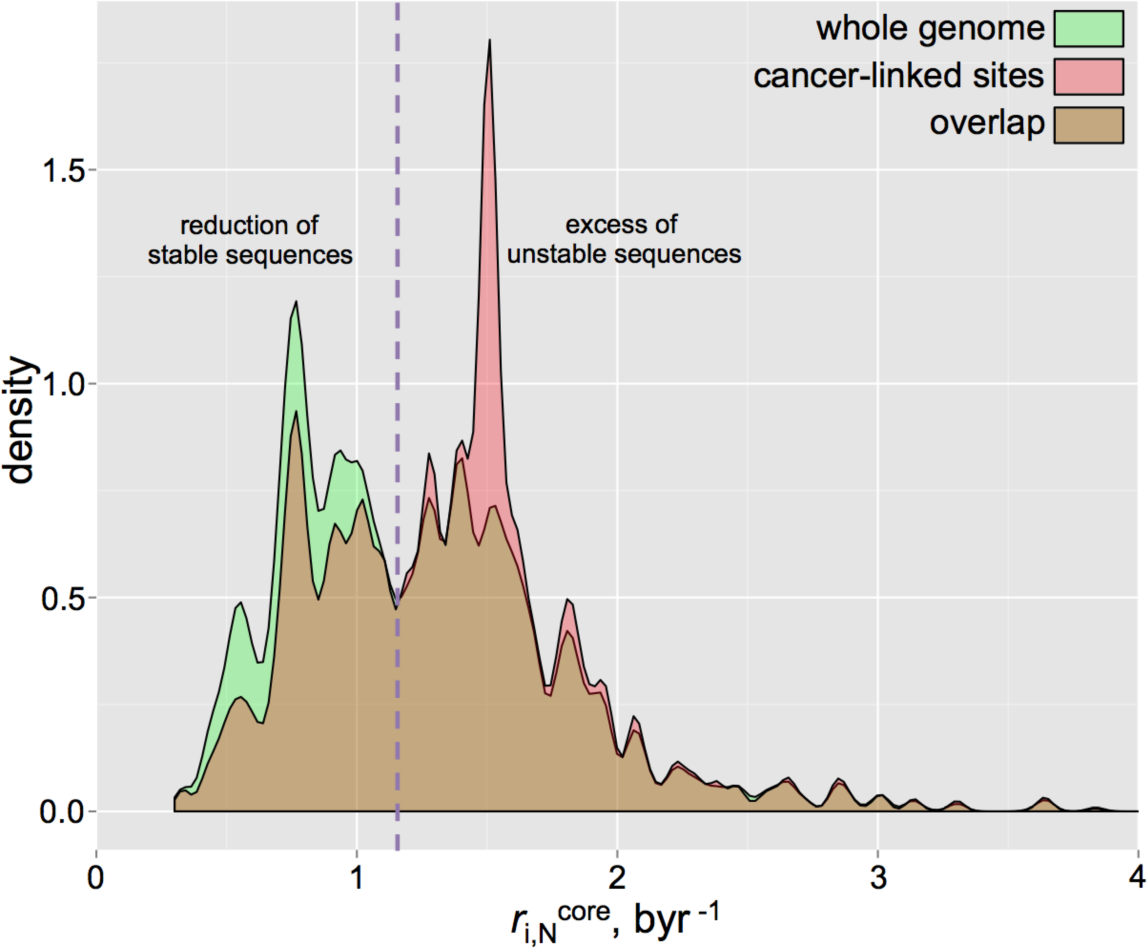
Basal substitution propensity (BSP) profiles of the whole human genome and cancer-linked somatic mutation sites. The density (kernel density estimate) distribution of the BSPs in the whole human genome (green), compared to the sites of the mutations associated with cancer (red) are shown. The overlaps of both distributions are in brown. The x-axis shows the BSP for the substitution to any other base 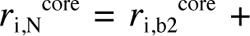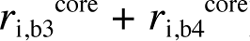. The comparison clearly shows a reduction of the stable sites and excess of the unstable sites in cancer-linked loci.

Like many other disease-causing mutation sites [56], most of the sites that are highly enriched in cancer (**fig. 6A**) are CpGs [57], which, even without accounting for the methylation driven increase [12-14] of the mutation rates, show high basal mutability [58]. However, **fig. 6B**,**C** demonstrates the discussed trend in the 7-mer cancer enrichment ratio (**Methods**) vs. BSP dependence even when all the CpG sites are removed from the analysis.

**Fig. 6.**
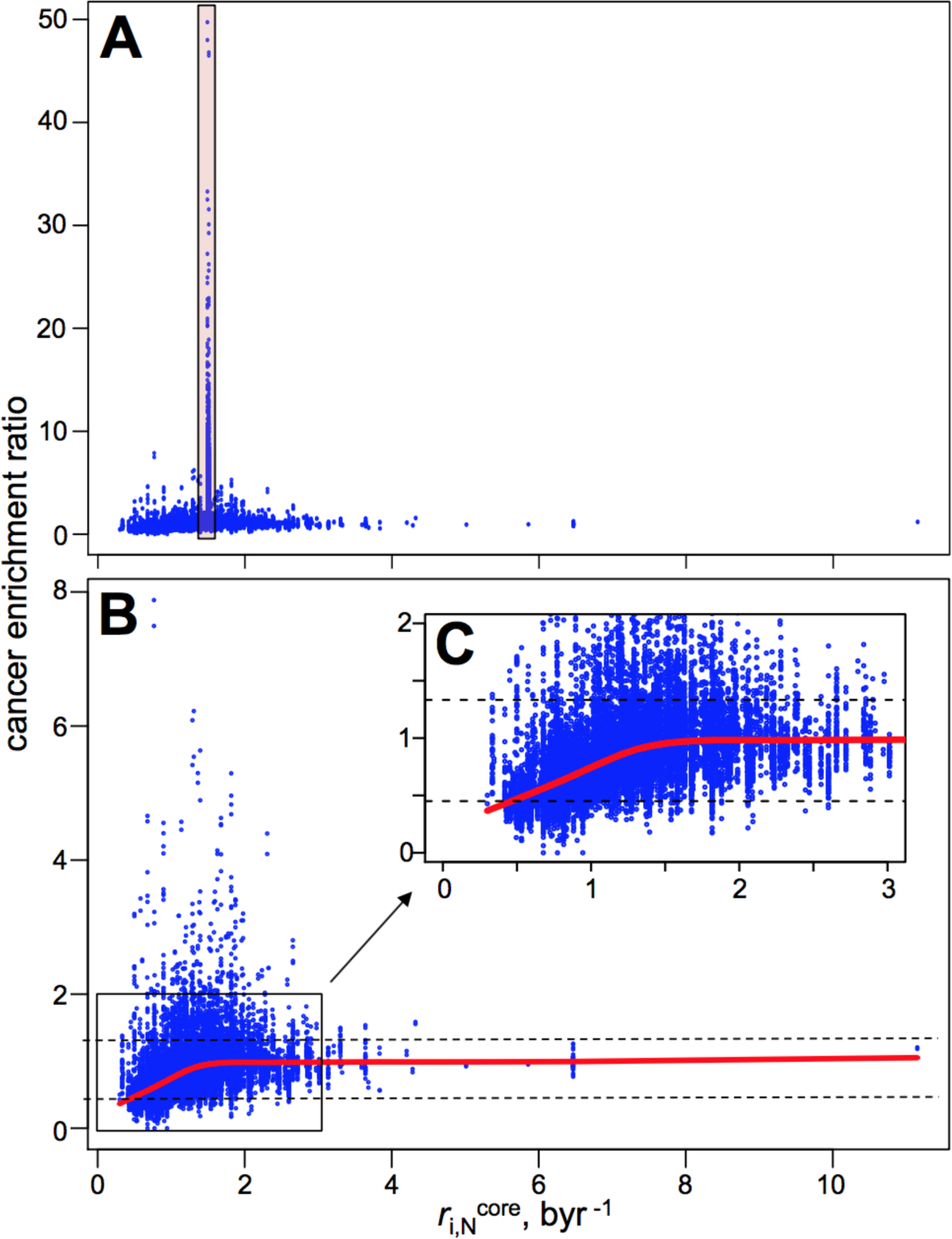
Enrichment of 7-mers with varying BSPs in the cancer-linked somatic mutation sites. The 4^7^ points in the plots correspond to unique 7-mer sequences. The cancer enrichment score for each such sequence was calculated by dividing the occurrence fraction of the sequence in only the cancer-linked sites to the fraction in the whole repeat-masked human genome. All the 7-mers that had either C or G of a CpG dyad at the centre show a remarkable cancer enrichment ratio (up to 49, see the points in the red box in **A**). Since for the CpGs, Trek data report on only the average C and G substitution rate constants, the BSP values for those points can be underestimated in **A**. However, even the average BSPs for C and G bases are higher than the discussed 1.13 byr^−1^ threshold. Accounting for the epigenetic methylation-driven increase in BSPs for the CpG containing sequences would only increase their BSP values. The plots **B** and **C** represent the data that exclude the sequences with CpGs at the centre. The mean cancer enrichment ratio for such subset was 0.89, with standard deviation of 0.44. The data points within the 0.89±0.44 range of cancer enrichment ratio are contained in between the dashed lines in **B** and **C**. The red lines in **B** and **C** represent the Lowess[65] fit, showing the decrease of the cancer enrichment ratio with the decrease in BSP.

Furthermore, while investigating the same relationship in different varieties of cancer (as classified based on the primary tissue and primary cancer types, see **Methods**) we can see that the trend is mostly in place for the 11 cancer types where we have enough data on non-coding somatic mutations (**fig. 7**), with the only exception being the oesophageal carcinoma (**fig. 7E**,**L**). The latter deviation might stem from the greater role of carcinogen driven mechanisms of somatic mutations in a tissue (oesophagal) more exposed to external cancerogenic agents.

**Fig. 7.**
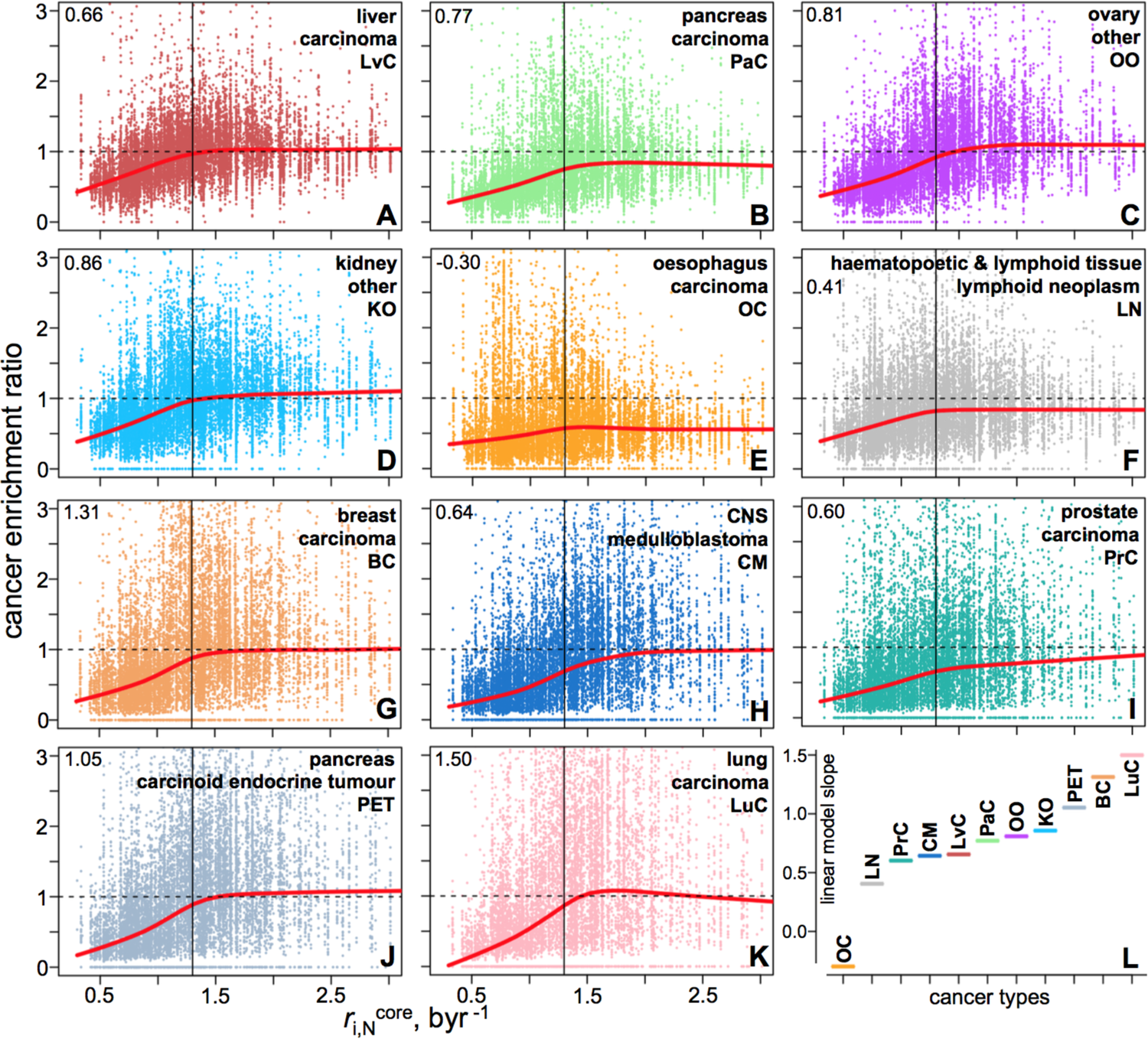
Enrichment of 7-mers with varying BSPs in somatic mutation sites linked to different cancer types. Each point in the plots **A-K** corresponds to a unique 7-mer sequence. All the 7-mers that had either C or G of a CpG dyad at the centre were excluded from the plots representing the zoomed [0, 3] byr^−1^ range of BSPs. The plots **A–K** show data from 11 cancer types, with their primary tissue and primary cancer types indicated at the top-right corner of each plot. The red lines in **A–K** represent the Lowess[65] smoothing fits, outlining the decrease of the cancer enrichment ratio with the decrease in BSP. The numbers at the top-left corners of the plots **A–K** show the slopes of the linear fits (not shown) for the data points below the 1.3 byr^−1^ (vertical line), where the depletion of stable k-mers is the most pronounced. The linear model slopes coming from all 11 cancer types are shown on **L** for comparing the extent of the cancer enrichment ratio vs. BSP dependence across the analysed cancer types.

Overall, our results show that the intrinsic BSP of different sites in DNA may contribute to their absence/presence in pathological genotypes. In particular, we observe that 7-mers with low germline BSPs of the central base are relatively depleted in cancer-linked somatic mutation data (**fig. 6 and fig. 7**). They present a cancer enrichment ratio that is smaller than 1, whereas for the unstable 7-mers, the enrichment ratio, on average, tends to 1, meaning that the presence in cancer is overall comparable to the one in the whole genome. These results outline the potential role of the general imbalance in proof-reading machinery in determining the accumulation of somatic mutations in cancer, where the mutations are originally caused by errors in replication that possibly emerge via mechanisms more or less common in somatic and germline cells.

## Conclusions

We have employed a single-genome and model-free approach (Trek) that reveals the core 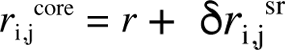 components of the spontaneous single-nucleotide substitution rates and basal substitution propensity constants (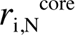) for the human nuclear genome. Although the mobile DNA elements have been used before [4, 39, 43, 59] for estimating averaged substitution rates, the increased quality of the human reference sequence and the detailed subfamily divergence studies for the L1 elements [35-38] done during the past decade enabled the construction of a specific direct method for the single-genome retrieval of the core 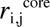 rate constants at a single-nucleotide resolution, while also accounting for the comprehensive short-range context effects beyond the previous Bayesian estimates for the +1/-1 base effects [34]. The retrieval of our 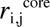 data in a single-genome manner adds additional value, since it ensures the absence of potential bias present in a) the comparison of the genomes of different species due to the differences in the molecular machinery (or presence of mutations in the respective genes) that influence the overall substitution rates, and b) in the SNP-counting based methods that rely on sites where the polymorphism can contain selection bias.

We used our context-dependent rate constants to provide a direct demonstration of the equilibration of a random DNA sequence into the one with overall (genomic base content) and short-range (different k-mer contents) characteristics that closely mirror that of the actual human genome. The L1-derived rates recapitulated fundamental properties of the repeat-masked and therefore L1-free human genome (**supplementary fig. S11**). This simulation thus provides significant evidence in support of the role of core neutral substitutions in shaping the compositional dynamics of complex genomes [60]. Importantly, our study demonstrates that the non-specific core substitution rates are capable of producing apparent selection or depletion patterns in the human genome. To this end, our *in silico* equilibrated sequences, obtained solely based on the full set of 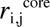 constants, can serve as better background standards for the comparisons to reveal real selection [61] for or against different sequence motifs.

Our calculated basal substitution propensities demonstrate a link between core substitution rates in the germline and the somatic mutations in cancer, outlining possible commonalities in the mechanisms of mutations at both levels.

The extended set of core substitution rate constants we report can potentially help advance our understanding in genome dynamics, with possible implications for the role of random substitutions in the emergence of pathological genotypes and the evolution of proteomes.

## Methods

The human reference genome sequence was taken from the Ensembl database (www.ensembl.org), and was of the version hg19/GRCh37. The positions and span of the retrotransposons were taken from the output of the RepeatMasker [62] processing, accessed through the UCSC genome database (www.genome.ucsc.edu). The repeat annotations were those corresponding to the version of the used human RefSeq genome (hg19/GRCh37). The R programming language (www.r-project.org) was used for all the consecutive analyses. Most of the computations were performed on the available Linux workstation and computing cluster facilities hosted at the Department of Chemistry, University of Cambridge, and the Cancer Research UK Cambridge Institute.

### Revealing the Core Substitution Rate Constants

All the remnant sequences of the mentioned subfamilies (**supplementary table S1**) were first aligned onto the 6064 nt reference sequence. As the reference, we took the consensus sequence of the human L1Hs retrotransposon (**fig. 1A**,**B** and **supplementary fig. S2**). The alignment was done in a pairwise manner with high end-gap penalties ("overlap" mode) that, while allowing insertions and deletions, did not severely break the queried sequences for false mappings with a better global alignment score. R with the Biostrings library for alignment was used. After the alignment, all the relevant substitution fractions were collected for each position in the five L1 subfamilies reporting on a specific time epoch (**fig. 1B**,**C**). For example, if the position *i* in the reference sequence was G(*b*_1_), the substitution rate constants were calculated for the G→A(*b*_1_→*b*_2_) transition and G→C(*b*_1_→*b*_3_), G→T(*b*_1_→*b*_4_) transversions. First, the base fractions were calculated for five time-reporting L1 subfamilies; i.e. to get the fraction of substitutions accumulated in ∼20.4 myr (age of L1PA5 [36]), all bases in L1PA5 remnants that were precisely mapped on the *i^th^* position of the reference sequence were counted and the fractions of G, A, C and T bases retrieved (**fig. 1C**). Here, we applied one of the robustness checks and made sure that the fractions were estimated if at least 700 mapped bases were present for the *i^th^* position in each time-reporting subfamily (**fig. 1B**). We also aimed to calculate such substitution rates for only the positions where the substitutions are random and not specifically selected for or against. In other words, the position should not be a polymorphic or a subfamily speciation-defining nucleotide. We filtered out such cases by ensuring that any eligible *i^th^* position had the same nucleotide of the reference sequence as its most prevalent variant with a minimum of 80% occurrence in all subfamilies (**fig. 1C**). The average crude single-nucleotide substitution rate is noted to be 12.85×10^−9^ substitutions per site per generation [5]. Assuming an average generation length of 20 years [2], the substitution rate constant in a time domain can be crudely approximated as 0.64 byr^−1^. In the course of 20.4 myr (the age of L1PA5), this should result in only a 1.31% substituted base fraction at a give site, caused by the average spontaneous substitution rates. Therefore, by assuming a threshold of 80%, we allow up to 15 times the variation of the rates from the average estimate, which is a safe range [5] for the direct estimation of the single-nucleotide substitution rates and their core variation. Having the substitution fraction data, from five different ages and for three (*b*_1_→*b*_2_, *b*_1_→*b*_3_, *b*_1_→*b*_4_) possible substitutions at the position *i*, allowed the fitting of a linear model via the least squares methodology for the fraction-versus-time dependence for each substitution separately (**fig. 1D**). If the data, hence the fitted line, were of high quality, the slope was expected to represent the 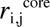 substitution rate constant. We applied the third robustness filtering at this stage, by making sure that the rates were calculated for only the cases where the time correlation of the substitution fractions in **fig. 1D** had greater than 0.7 Pearson’s correlation coefficient. This ensured that the retrieved fractions of the substitutions comprised of only the time-accumulated substitutions, rather than of targeted substitutions during the active life-span of the L1 elements, before their silencing. Please note, however, that the correlation coefficients in most of such time correlations that passed the whole Trek procedure were substantially higher (the observed Pearson’s correlation coefficients were centred at 0.92 with 0.07 standard deviation). Furthermore, the resulting slope (rate constant) estimates had significantly high t-values, averaged at 6.2, showing that the standard error in estimates was, on average, 6.2 times smaller than the estimated value. The individual t-values are presented in the **supplementary data** along with the rate constants. The procedure was done for all the 6064 positions in the L1 reference sequence, except the positions 5856–5895 and 6018-6064, close to the 3’-end (**supplementary fig. S2**) that engulf low-complexity G-rich and A-rich sequences correspondingly, prone to alignment errors. One of the reasons for the usage of only the young L1 subfamilies (spanning 20.4 myr age) was to minimise the potential error in rate constant determination in the Trek procedure caused by repeated substitutions hitting the same position during the considered period of the substitution accumulation. The effect is indeed negligible for 20.4 myr span, as we can estimate using the above mentioned 0.64 byr^−1^ value [5] for the average *i→j* substitution rate constant, *r*. Since the rate constant is sufficiently small to induce only a small δ*f_j_* change in substituted base fraction during the δt = 0.0204 byr (20.4 myr) time period (see above), we can equate the δ*f_j_* change in the fraction of the base *j* (at the given position that had the original base *i* identity in a large population of homologous sequences) to the p≈δ*f_j_* ≈ rδt substitution probability within *δt* period. We can thus make a crude estimation for the probability of the second substitution to another, *k*≠*j*, base happening at the same position to be (*r*δ*t*)^2^, which is the product of individual substitution probabilities assuming that the rate constant does not change from our average estimate *r* across those two substitution types. We can permit this for the sake of the back-of-the-envelope estimation of the order of the effect expected from the repeated substitutions hitting the same site within 20.4 myr period. To this end, the δ*f_j_^app^* apparent change in *i*→*j* substitution fraction that we would observe by neglecting the additional *j*→*k* substitution, would underestimate the more realistic δ*f_j_* and be equal to δ*f_j_^app^* = rδt-(rδt)^2^, as we would not count the *j* bases that emerged but became additionally substituted by *k*. Hence, the corresponding apparent rate constant that neglects second substitution would also underestimate the actual value *r*, and can be expressed as *r^app^* = δ*f_j_^app^*/δt = (*r*δt - (*r*δ*t*)^2^)/δt=*r*(1-*r*δt). This means that the underestimation of the actual rate constant would be by [*r*-*r*(1-*r*δt)]×100/r = 100×rδt%. Putting 0.64 byr^−1^ for *r* and the 0.0204 byr for δt, we can expect only 1.3% contribution from the repeated second substitution at the same position within 20.4 myr. Since some of the other non-*j* bases could become *j*s and balance the underestimation of the δ*f_j_* fraction, the error could be even smaller. This shows that repeated substitutions could be safely ignored in 20.4 myr time-scale. Furthermore, the validity of our 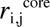 rate constants was further checked through the two independent analyses reflected in **supplementary fig. S3** and **fig. S4**.

### Finding the Influence Range of Neighbour Nucleotides

We have used generalised boosted models [63] (GBM) to elucidate the effective range for the core sequence-context effects. This was achieved by developing test models to evaluate the predictive strength of only the neighbouring bases in defining the core substitution rate of the central base. The GBM is a machine learning methodology that produces a regression or classification model by assembling weaker models in the forms of decision trees. It was used as implemented in the gbm library for R. For each *i*→*j* substitution type, all the found Trek data were taken without the possible outliers, which were filtered by allowing only the usage of the values that were within the 1.65 times standard deviation range (keeps ∼90% data if normally distributed) of the constants in a given substitution category. The sequences were then processed to produce *pos/b_i_* uncoupled features that were associated with the relative adjacent positions (*pos*, – for upstream and + for downstream positions) and their possible four *b_i_* base types. Those features took values 0 or 1, depending on whether the base at an associated relative position was of the *b_i_* (1) or any other base type (0). For instance, if we want to develop a model accounting for only a single upstream (*pos* = -1) and a single downstream (*pos* = +1) nucleotides, hence predicting the substitution rates for different 3-mers, where the central base is the one that mutates, then we would produce 8 *pos/b_i_* features for the GBM fitting. There, 4 binary features (-1/A, -1/C, -1/G and -1/T) would describe whether the upstream -1 position is of base type A, C, G or T, and 4 binary features (*pos/b_i_*) would describe the same for the downstream +1 position. We built the models using 3-, 5-, 7-, 9-, and 11-mers, thus accounting for 1, 2, 3, 4 and 5 upstream and the same number of downstream neighbour bases. The absence of the coupling in the binary features, unlike in the case where, for instance, one employs only two binary features per 4 states, enabled us to also investigate the predictive significance of each nucleobase identity at a given neighbouring position, which was useful in deciding against the construction of more complex machine learning models (see below) using additional features with higher level of abstraction for the sequence information (overall base content, sequence-derivative properties). The GBM models were then fitted by systematically trying different permutations of the tuning values [63] for the number of trees (50–7500), interaction depth (1-10), shrinkage (0.001, 0.01 and 0.1), the number of minimum observations per node (1-28) and the bag fraction (0.25-0.65). The optimal combinations of the tuning parameters were found per substitution type and sequence length, via a 16-fold cross validation repeated 7 times. The found best parameters are accessible in the **supplementary data**, and the predictive performances of the best models, from the repeated cross validation studies, are presented in **supplementary fig. S5**. To make a predictor of 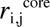 based on a sequence only, we found it much better to use direct values coming from the proposed Trek methodology, rather than the GBM models, as the Trek values are already well averaged across multiple occurrences of the same sequence in different loci of the human genome (**fig. 1-3** and **supplementary fig. S3**). Furthermore, the overall poor performance of the GBM models implies that the influence of the immediate context is highly non-additive and non-Bayesian, which is expected taking into account the nature of the core context-dependent substitution rates. The latter rates reflect the intrinsic short-range sequence properties, interactions and recognition with the overall mutagenic and repair machinery present in a given organism. There, the whole sequence at a certain small scale [9] is what defines the interaction [11], and it is hard to represent such effects through even smaller-scale constituents. The direct model-free approach used in our Trek mapper methodology (see below) thus seems essential in mapping the 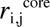 rate constants throughout the human genome. To this end, the GBM models here had a sole purpose of identifying the optimal range of influence for accounting the neighbouring nucleotides. The optimal range was found to be captured, on average, by a 5-7-nt long window (**supplementary fig. S4** and **S5**) which is in an excellent agreement with the prior <10 nt estimate [9-11, 45, 46]. We thus used the maximum 7-nt length to stratify the Trek data for the further model-free mapping on any provided sequence, including the whole human genome.

### Mapping the Trek Substitution Data on any Sequence

We developed a Trek mapper program. For each *i* position in a query sequence (**supplementary fig. S6**), the program looks at the bases *i*-3 to *i*+3. If the exact 7-mer, with the associated rate constant values, is not available in the Trek database, the program reduces the size of the sequence to 5, by considering *i*-2 to *i*+2 positions, or, if necessary, to 3- or 1-mers, until an exact match is found in the database. This would essentially mean that some reported substitution rates would come from the actual triad data. About half will come from pentads and some from heptads, accounting for more precise sequence-context information. A few will originate from the fully averaged single-base (1-mer) Trek rates. Single-base values are also returned for the terminal positions in a query sequence. For each unique sequence in the discussed 7-, 5-, 3- and 1-mers, if the k-mer appears more than once in the reference L1 sequence, of course with different neighbours at the positions out of the k-mer range, we average the Trek values by taking the median. For instance, the *r*_G→A_ substitution rate constant in the 3-mer AGT represents the rate averaged across all the appearances of AGT in the L1 reference sequence, which would normally be with varying other neighbour bases, out of the 3-mer range. The *r*_G→A_ in AGT will therefore represent the average rate constant across all the representatives of the significant range, the NNAGTNN 7-mers, present in L1, where N can be any of the four bases. In the same way, the substitution rate constants for the single bases (1-mers) can be considered as fully averaged across all the possible neighbour effects in NNNGNNN sequences. Our algorithm therefore makes the most of transposon exposed substitution rate data of the human genome, returning the best possible values inferable from our Trek database and, where uncertainty is present, returning the best averaged values for a shorter context range. Furthermore, we have enabled the usage of symmetrised Trek parameterisation, assuming an overall strand-invariance of the substitution rates. In the latter case, the complementary rate constants of the central bases in two reverse complementary k-mers were equalised. For example, the G→A substitution rate constant (*r*_1_) in the 3-mer AGC was set equal to the complementary C→T rate constant (*r*_2_) in the reverse complementary GCT. The data equalisation was done in the following way: if both *r*_1_ and *r*_2_ were of the same quality accounting for the whole sequence-context information in both 3-mer variants, then both values were set to (*r*_1_+*r*_2_)/2; however, if one of the rate constants was determined with a better quality, since the full 3-mer data for the other case was missing and the 1-mer average was used as a replacement, then the rate constant of the better quality variant was assigned to both *r*_1_ and *r*_2_. Accounting for the strand symmetry improves the results of the validation studies, further refining the substitution rate constant values and increasing the coverage of longer k-mers in the Trek database.

The described Trek mapper program, along with the associated data can be accessed through the http://trek.atgcdynamics.org web page. Future improvements in data and the program, through extending the types of mobile DNA in the Trek procedure, will be reflected on the same web site. The Trek mapper server application was written in R, using the Shiny library and server backend (http://shiny.rstudio.com). The open-source stand-alone program, supporting both graphical and programmatic (terminal) interaction and multi-processor computing, can be obtained from the same web page, to be used for larger projects and genomes.

### Equilibration of a Random DNA Sequence with the Trek Rates

A 5 million (mln) nt sized random genome was created with the initial A, T, G and C base contents set to 20, 20, 30 and 30% correspondingly, hence with 60% genomic G+C content. The length was selected to cope with the finite computational and time resources, though operating on lengthier sequences will not change the outcome of the calculations, since the captured sequence-context effects are within 7-nt window. We first calculated the probabilities of all the possible substitution in this random sequence, which basically meant the assignment of 3 substitution rate constants per position in the genome, describing the conversion into the three bases other than the base already present in the respective position. This was done using the Trek mapper described above. Next, at each step, we sampled 5000 substitution weighted by the calculated 3×5 mln rate constants. We then identified those 5000 positions and the corresponding substitution types that were sampled to happen (**supplementary fig. S7**), performed those substitution, and, updated the probability values via the Trek mapper. Repeated multiple times, the process evolved the sequence (**supplementary video 1**) ruled by the core spontaneous substitution rate constants that are sensitive to the changes in the sequence composition at the immediate vicinity in the genome.

For the comparison of the simulated sequence at equilibrium with the real human genome (RefSeq), we calculated the fractions of different oligomers (k-mers) in both sequences (**supplementary data**). The k-mer contents of the human genome were calculated by sliding a window of size k (from 1 to 7) and counting the occurrence of each 4^k^ unique sequence. We used a direct calculation of the lexicological index [64] of a string to increase the computational efficiency of the k-mer counting. Although, data from the masked human genome were used in the k-mer analyses to rule out any bias from the presence of the same L1 elements in the object of application of Trek data, the comparison of the masked and unmasked genomes showed only negligible differences in both single base and short k-mer contents. If, however, we consider only the L1 elements, the k-mer content is quite different from the rest of the genome.

### Basal Substitution Propensity Constants at Cancer-Linked Sites

We took all the non-coding somatic point mutation data associated with cancer from the COSMIC database [55] (www.sanger.ac.uk/cosmic, NCV dataset accessed in February, 2015). Since our Trek rate constants are for the spontaneous core substitutions, we only considered the sites that were also not declared as known SNPs (the status was present in the NCV dataset). This was to ensure that we excluded sites where an active polymorphism is potentially encouraged by natural selection. About 3.7% data from the remaining set of cancer-linked somatic mutations were duplicates, with no differences found in genomic location and mutation types. We removed those, keeping only the single first-encountered copies of such entries. The resulting data contained 5984711 mutation entries.

The cancer enrichment score (**fig. 6**) for a given k-mer sequence was calculated by taking the ratio of the occurrence fractions, calculated in (numerator) only the cancer-linked sites (where the linked base is the central one in the k-mer) and (denominator) in the whole repeat-masked human genome (**supplementary data**).

We next repeated the above analysis by examining non-coding somatic mutations from different types of cancer, as guided by the primary tissue (primT) and primary caner (primC) types recorded in the COSMIC database. To stratify data, we examined the primT:primC pair as a cancer type identifier for each mutation. The data contained 29 unique primT:primC pairs, of which 11 had a substantial number of records (2422060 non-coding mutations for liver:carcinoma, 1287384 for pancreas:carcinoma, 851028 for ovary:other, 411076 for kidney:other, 331520 for oesophagus:carcinoma, 286800 for haematopoetic and lymphoid tissue:lymphoid neoplasm, 92276 for breast:carcinoma, 79342 for central nervous system (CNS):primitive neuroectodermal tumour/medulloblastoma, 69721 for prostate:carcinoma, 62714 for pancreas:carcinoid endocrine/tumour, 38835 for lung carcinoma). This list was followed by 19004 records with non-specified primary tissue and cancer types, and substantially low number of records for the rest (at or below ∼10k records). We thus analysed the results from the top 11 identifiers with the most number of recorded somatic mutations.

## Supplementary Material

The Trek 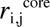 database and the mapper are housed at http://trek.atgcdynamics.org. Additional figures and a table referenced in the text (**figures S1-S11, table S1**), as well as the detailed descriptions of the other supplementary files, can be found in the **supplementary information** pdf file. We deposit a mov format video showing the *in silico* dynamics of a random sequence upon equilibration (**supplementary video 1**). The raw data on the substitution rate constants in the reference L1 sequence (**data 1**), the resulting Trek database processed with (**data 2**) and without (**data 3**) the strand-symmetry considerations, the k-mer content for the masked and unmasked human genomes (**data 4**), the full set of 7-mer sequences with the respective cancer enrichment scores and basal substitution propensity values (**data 5**), and the GBM parameters that were minimising the error of the tree-based test models (**data 6**) can all be found in the single **supplementary data** text file, under the corresponding sections. All the associated method and analyses source codes are available from the authors upon request.

## SUPPLEMENTARY INFORMATION

**Table S1.** Number of genomic copies and estimated age of hominoid L1 subfamilies. The numbers of L1 mobile elements (N^hg^) in the human genome were revealed through the RepeatMasker (Smit et al. 2015) processing of the genome. The divergence age estimation was obtained from the published molecular clock analysis (Khan 2006).

| **L1 type** | **N<sup>hg</sup>** | **age, myr** |
| --- | --- | --- |
| L1Hs | 1528 | 3.1 |
| L1PA2 | 4867 | 7.6 |
| L1PA3 | 10565 | 12.5 |
| L1PA4 | 11763 | 18.0 |
| L1PA5 | 11171 | 20.4 |

**Figure S1.**
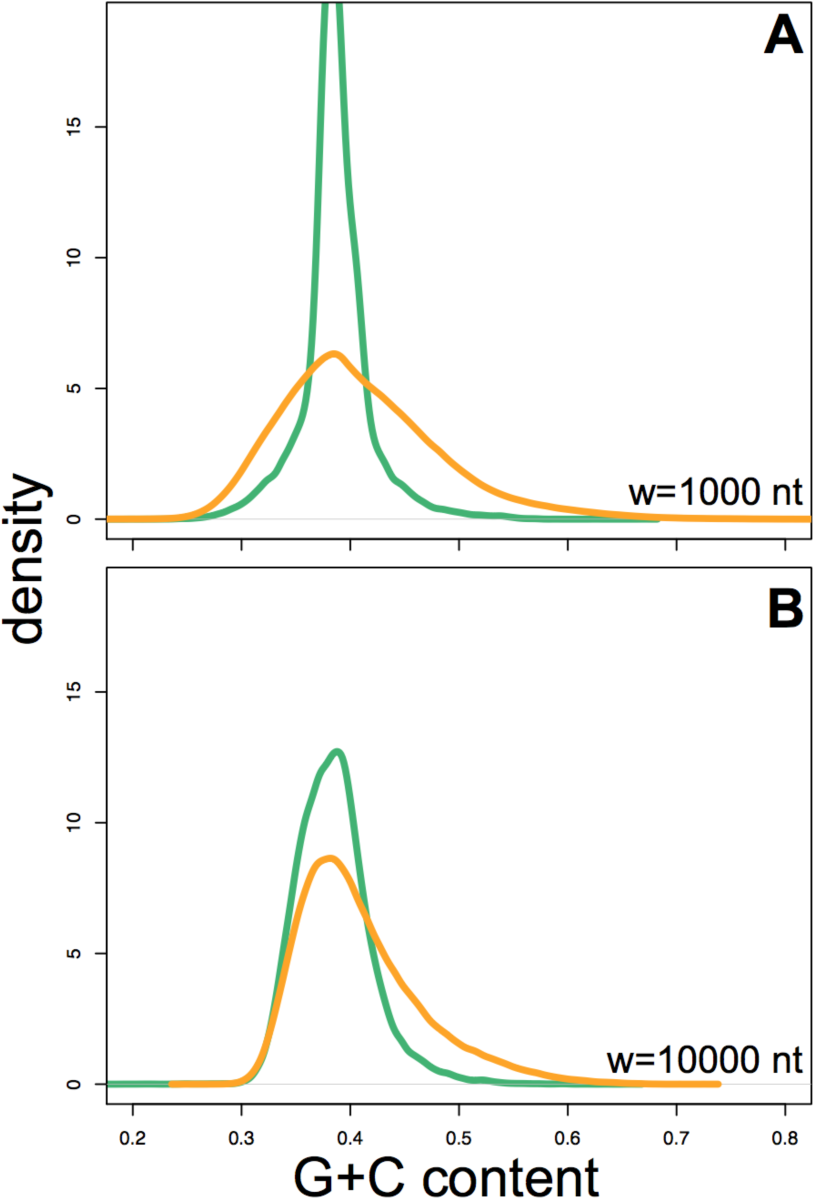
Long-range G+C context of the L1 insertion sites in the human genome. The distribution of the G+C contents for all the w-sized (1000-nt in **A** and 10000-nt in **B**) bins in the human genome (orange lines) is shown, as compared to same distribution but using only the bins centred at the midpoints of all the remnants of young L1 elements (L1Hs, L1PA2, L1PA3, L1PA4, L1PA5).

**Figure S2.**
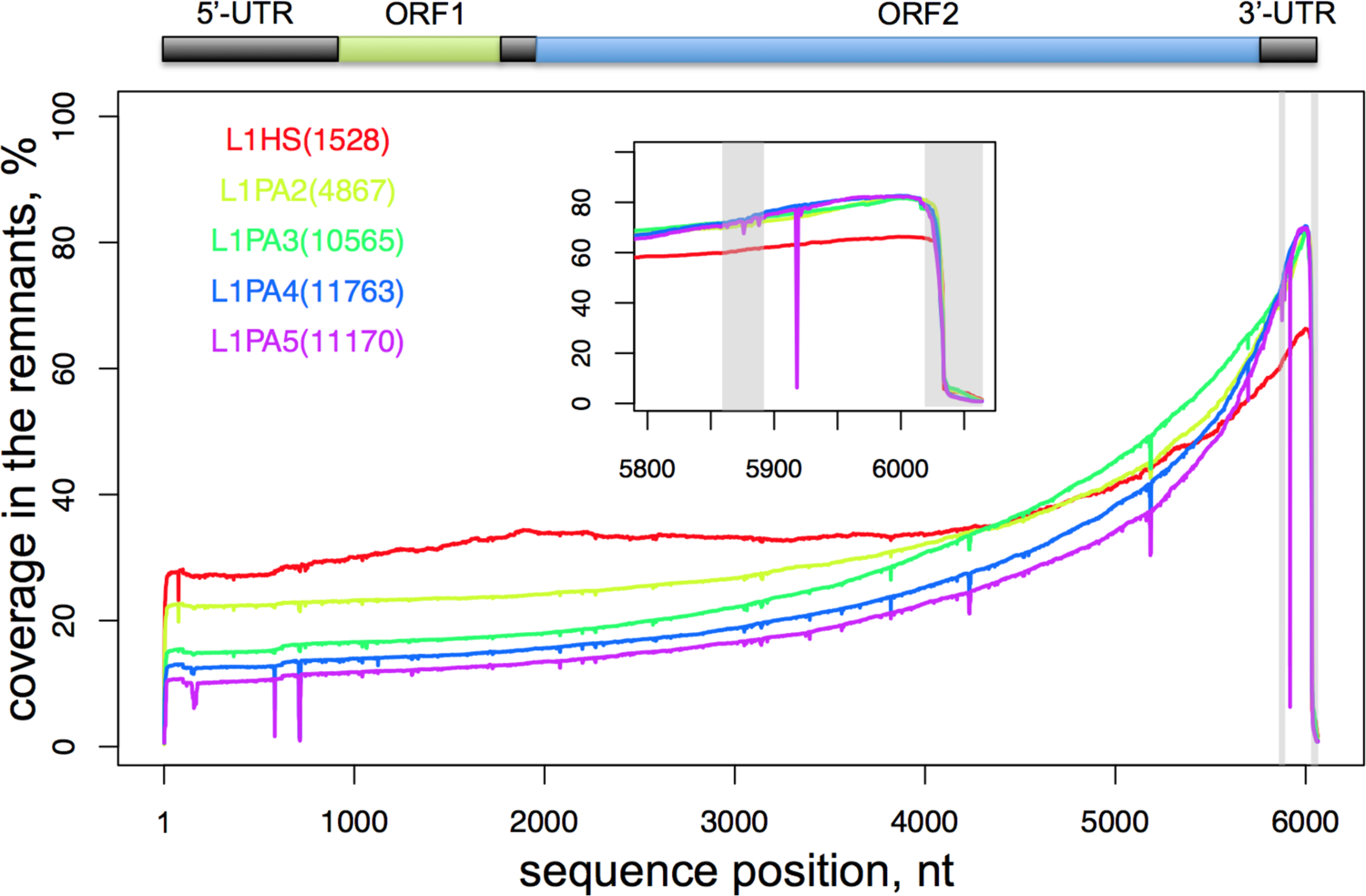
The position-wise coverage of the five subfamilies of L1 remnants in the human genome. All retrotransposon types, which are not too ancient for the time-accumulated substitutions to severely decrease the information content, were pairwise aligned on the 6064-nt consensus sequence of the human-specific L1Hs subfamily (reference sequence). The graph shows the percentage of cases where, for each considered L1 subfamily, the remnant sequences were mapped onto the corresponding position (x-axis). The gene organisation in these L1 elements is displayed on top, highlighting the terminal untranslated regions, along with two open reading frames (ORF1 and ORF2) and the short inter-ORF region. The positions 5856-5895 and 6018-6064, close to the 3’-end (also zoomed in the sub-plot) that engulf the low-complexity G-rich and A-rich sequences are marked with grey bands and excluded from the substitution rate analyses. The colour coding of the examined hominoid L1 retrotransposons, along with the genomic copy numbers in the human genome, is shown on the plot. The abrupt drops in the coverage at different positions along the sequence are because of deletions. The figure highlights the incomplete 5’-reverse transcription, characteristic to LINE elements. The poly-A tail at the 3’-end is also largely incomplete in most L1 remnants. It is interesting to note that the further back in time we go in terms of the activity period of the L1 subfamily, the less preserved the 5’-side of the L1 elements become relative to the 3’-end. An overall decrease of the preservation, hence coverage, is expected for the more ancient subfamilies, but the observed decrease relative to the 3’-end can indicate that the more recent subfamilies evolved a more efficient retrotransposition and/or improved transposonic RNA stability that results in more complete insertions.

**Figure S3.**
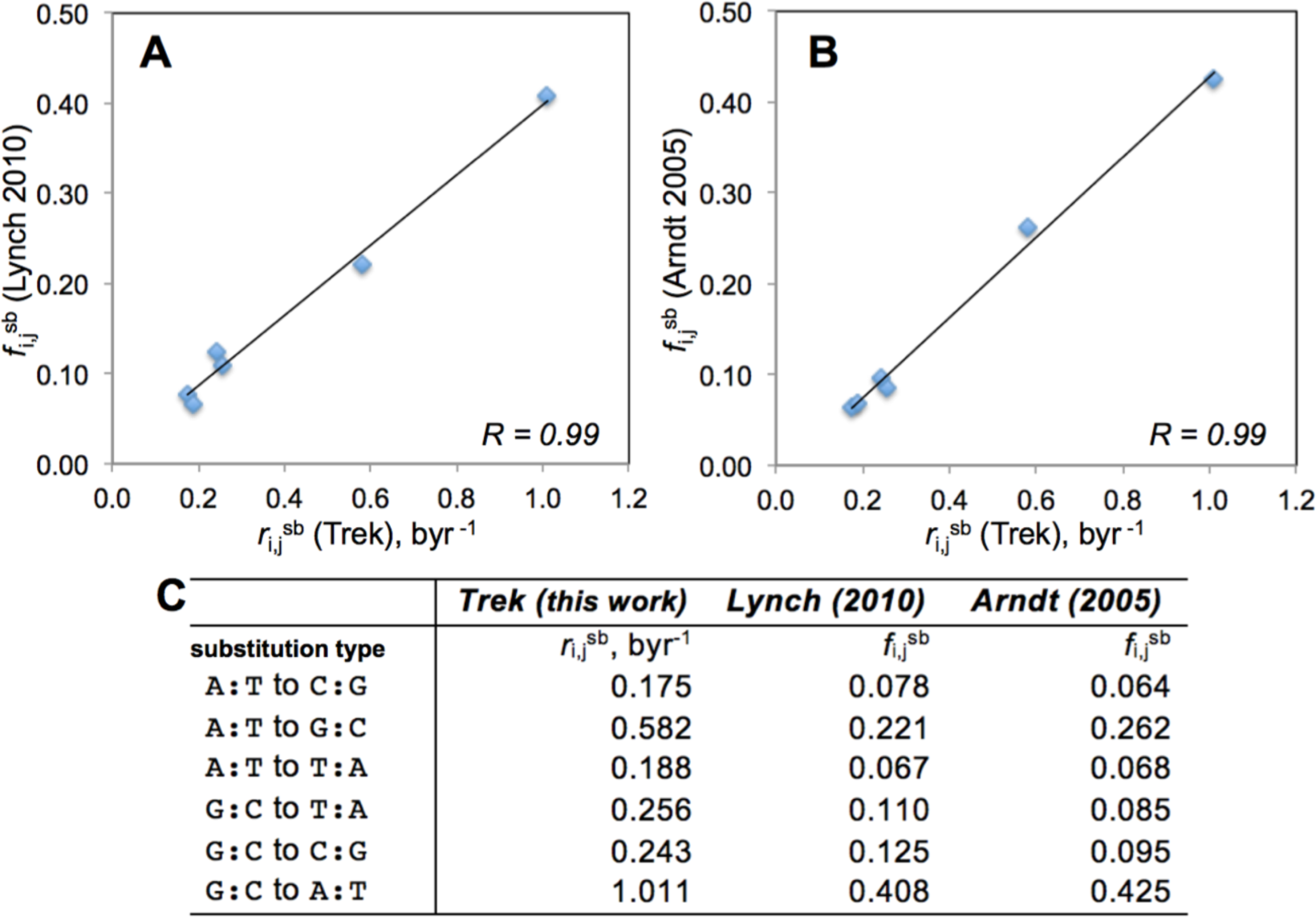
Demonstration of the unbiased averaging in the Trek substitution rates. The median values of the Trek-reported 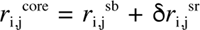 substitution rates, which should be reasonable estimates of single-base genomic average 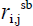 rates (in time domain), are compared (**A, B**) with two published datasets (Arndt et al. 2005; Lynch 2010) for 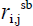, expressed by normalised substitution fractions. Since both datasets report on strand-symmetry-accounted six unique rates, we have performed the same strand-symmetry averaging of the median values shown in **fig. 2** before the comparison. The numerical data are presented in **C**. The Pearson’s correlation coefficients are shown on the plots **A** and **B**.

**Figure S4.**
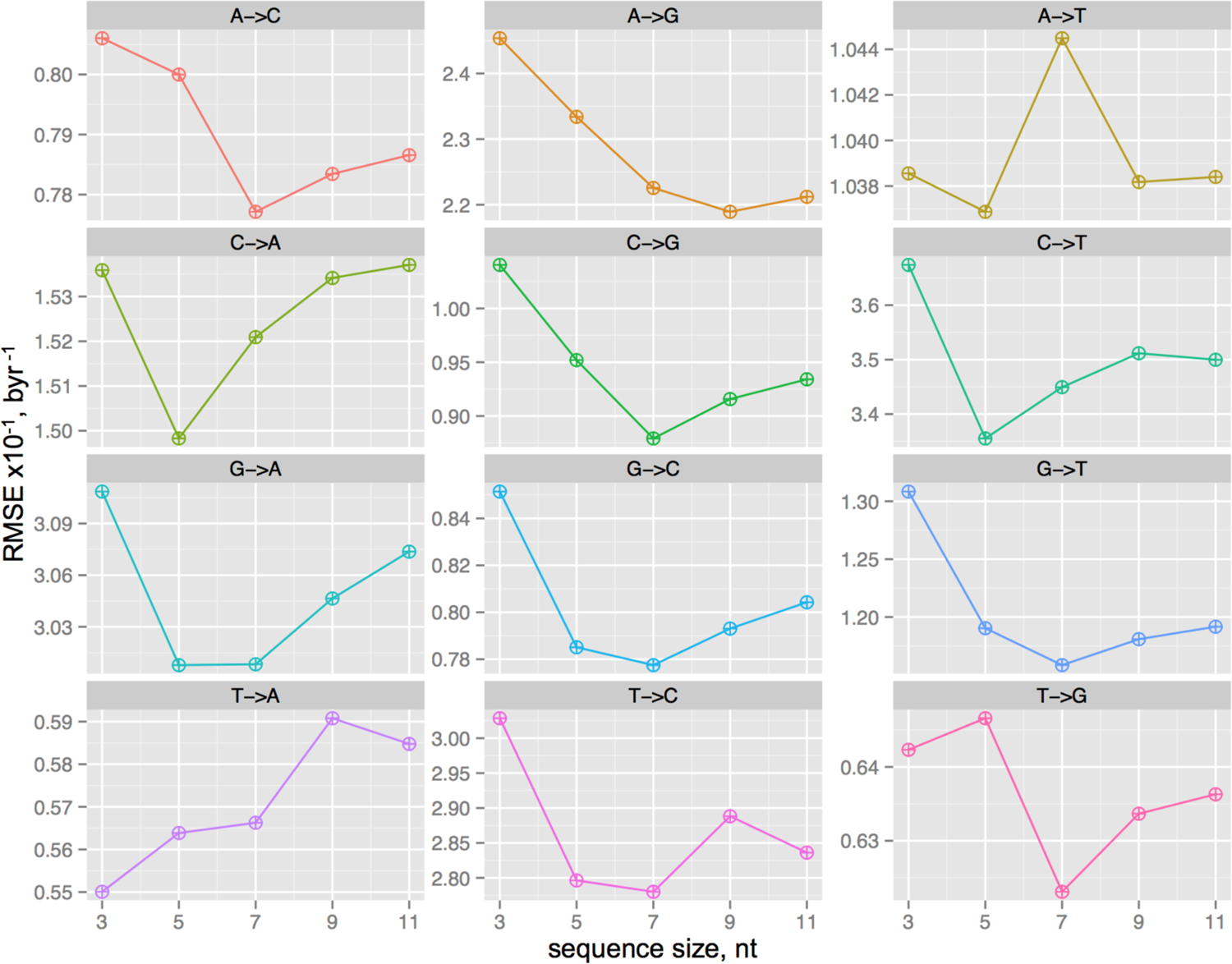
Selection of the optimal short-range sequence length directly influencing the substitution rates. The root-mean-squared errors of the rate constant prediction from only the neighbouring base information is presented as a function of the accounted sequence length, where the substitutions occur at the central base. The examined lengths are thus odd numbers, to allow equal number of upstream and downstream bases around the substitution point. The test models for the rates, to assess the optimal sequence length, were built using the machine learning, generalised boosted models. The 5- and 7-nt sequence lengths are found to be the best for general applicability for all the substitution types. Note that the machine learning predictions here are solely for finding out the optimal sequence length and, for the actual 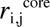 determination, a direct model-free mapping to the Trek substitution database is used.

**Figure S5.**
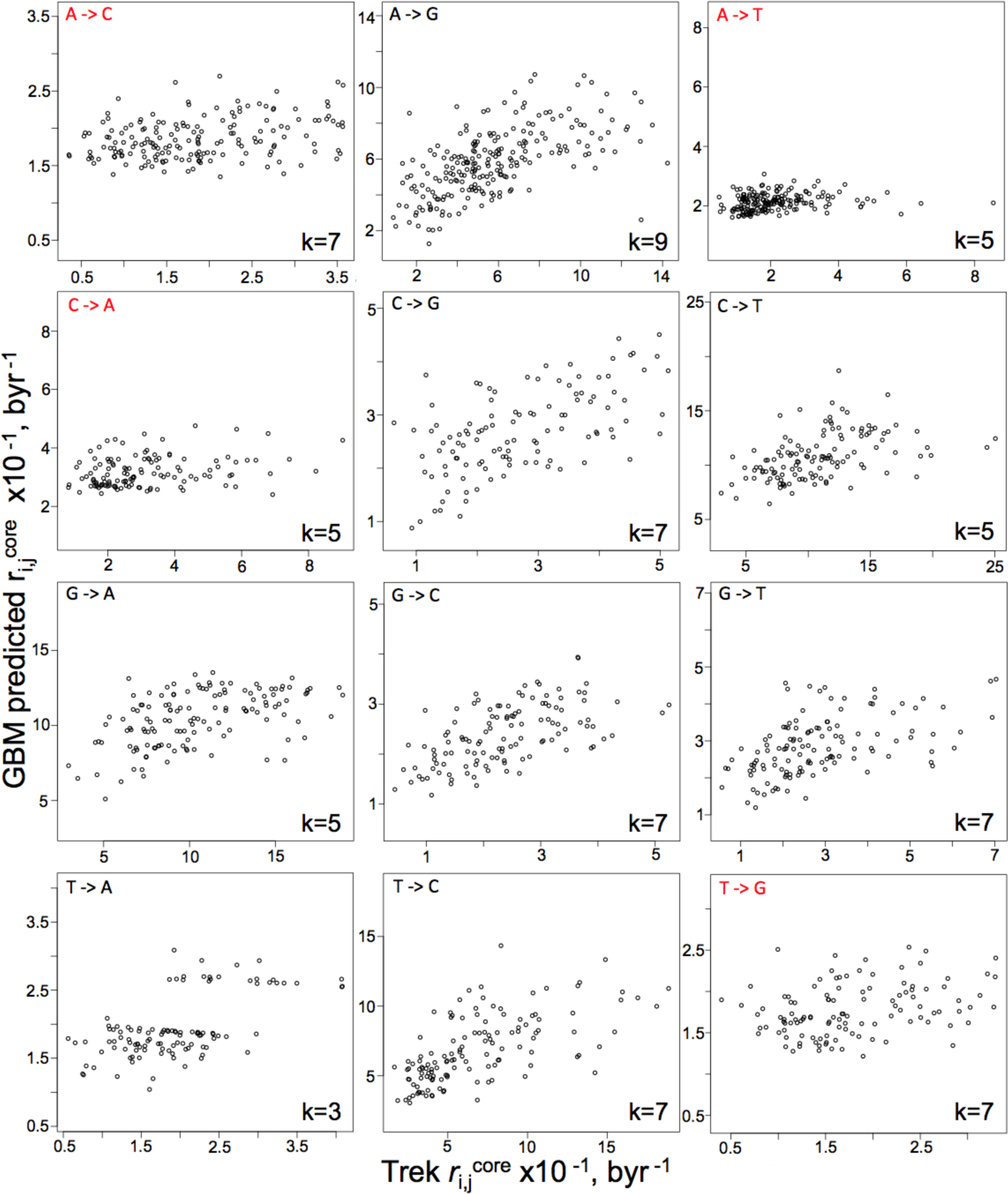
Performance of the optimal GBM models in our test constructs. Machine learning models were built to predict the 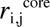 constants using only the knowledge of the neighbouring bases. The resulting highest performing lengths of the sequences (k-mers) are shown at the bottom right corners of each plot. The best models identified for all *i→j* substitution types are presented here as an example of the predictability of the substitution rates from the neighbouring residues. We have used these analyses to infer the optimal sequence (k-mer) length, found to be around 5-7 nt. We then utilised the found maximum length (7 nt) for all the substitution types as a factor to stratify the Trek 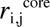 constants for the direct mapping to any given sequence.

**Figure S6.**
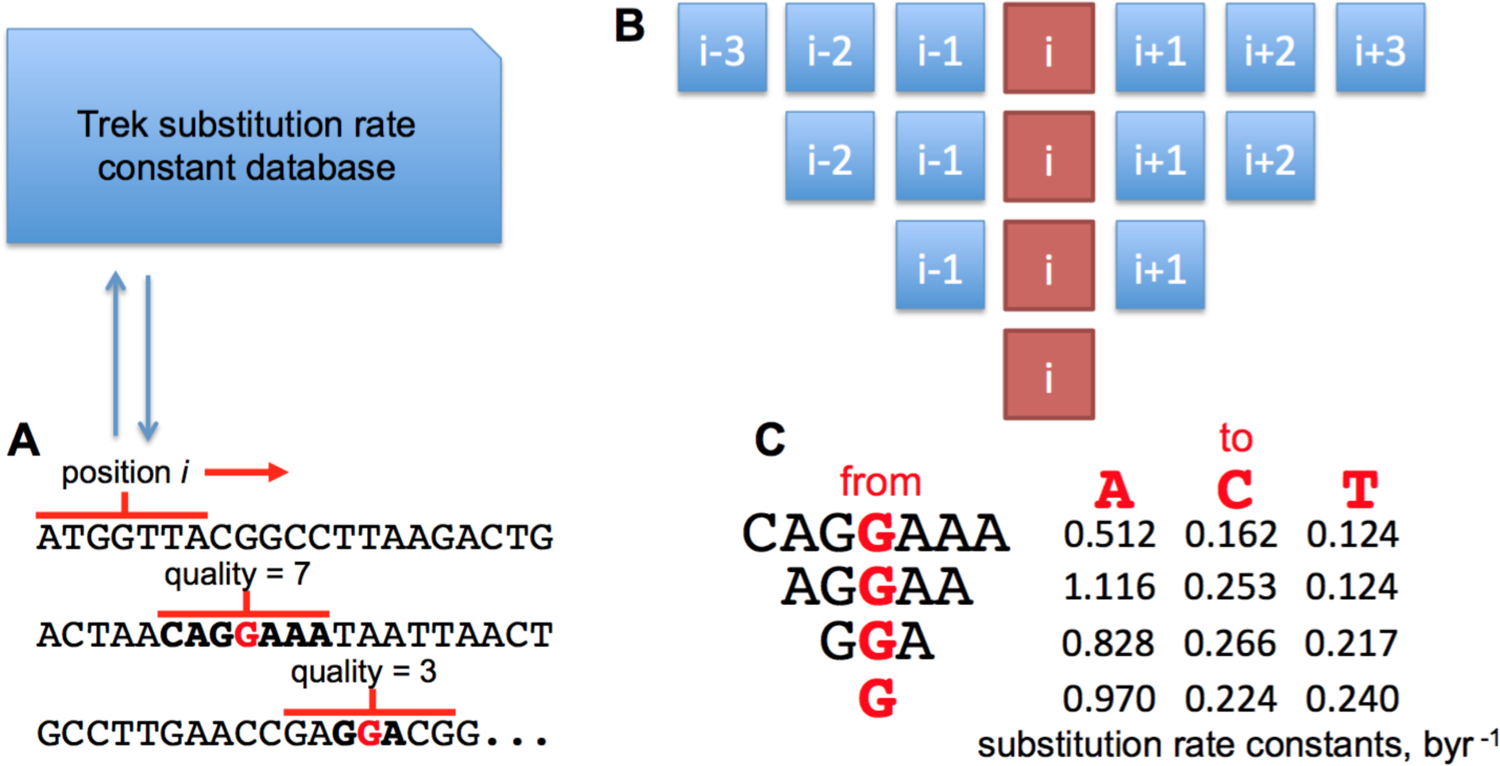
The procedure of mapping the 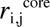 constants onto any sequence. A query sequence (**A**) is analysed by examining each base via a 7-nt window centred at the position of the base. To reveal the rate constants for the three substitutions of the central base into the three other bases, the Trek database is searched. The Trek data contain information on the full set of unique k-mers (k = 1, 3, 5 and 7) found in the reference L1 element, along with their respective three substitution rate constants. The Trek data are averaged where multiple values are found because of multiple k-mer copies in the same L1 reference sequence. In case a representative match is not found with the full 7-nt long sequence, the window around the given position in a query sequence is shortened (**B**) into the longest variant (5-nt, 3-nt or 1-nt) with a match detected in the Trek database. **C** demonstrates a case for the average substitution rate constants of the base G mutating to A, C or T, with different extent of context information in the Trek data.

**Figure S7.**
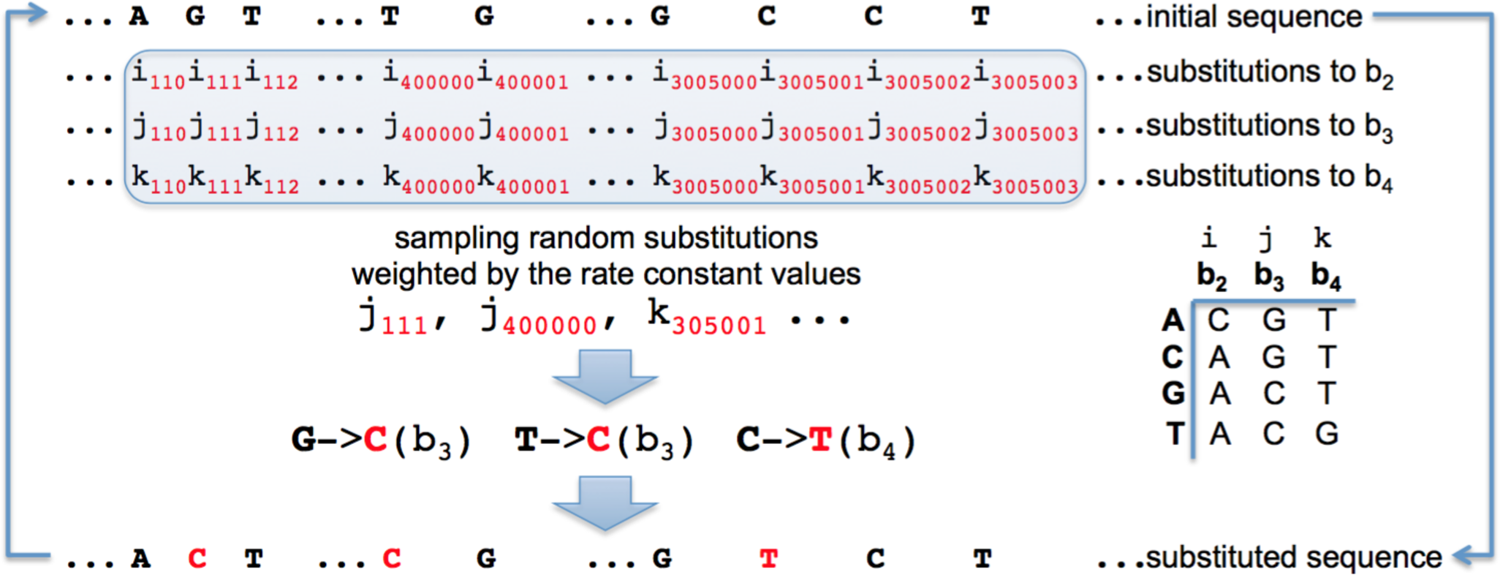
The *in silico* evolution of a genome with only 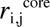 substitution rate constants. A random sequence of 5-mln-nt is generated with 60% G+C content (30% G, 30% C, 20% A, 20% T). Next, for each position, *pos*, the *i_pos_*, *j_pos_* and *k_pos_* rate constants for the substitutions into the other three b_2_, b_3_ and b_4_ bases are obtained from the Trek database, using the 7-nt sequence window centred at each *pos*. This generates 15-mln (3 times the sequence size) rate constant values that, for the first-order kinetic processes, are proportional to the substitution probabilities. We then sampled 5000 instances from the 15 mln generated set of *i, j* and *k*, with the sampling process weighted by the *i_pos_*, *j_pos_* and *k_pos_* values. Those sampled instances contain information on both the substitution positions (*pos* subscript) and the substitution types (*i, j* and *k* describing the substitutions to b_2_, b_3_ and b_4_ bases respectively, the latter ones being bases other than the initial base, always ordered alphabetically). After performing the sampled substitutions, we repeat the cycle and continue the process each time with substitution rates updated wherever the already performed substitutions affect the values. We continue the simulation until the equilibration of the sequence composition.

**Figure S8.**
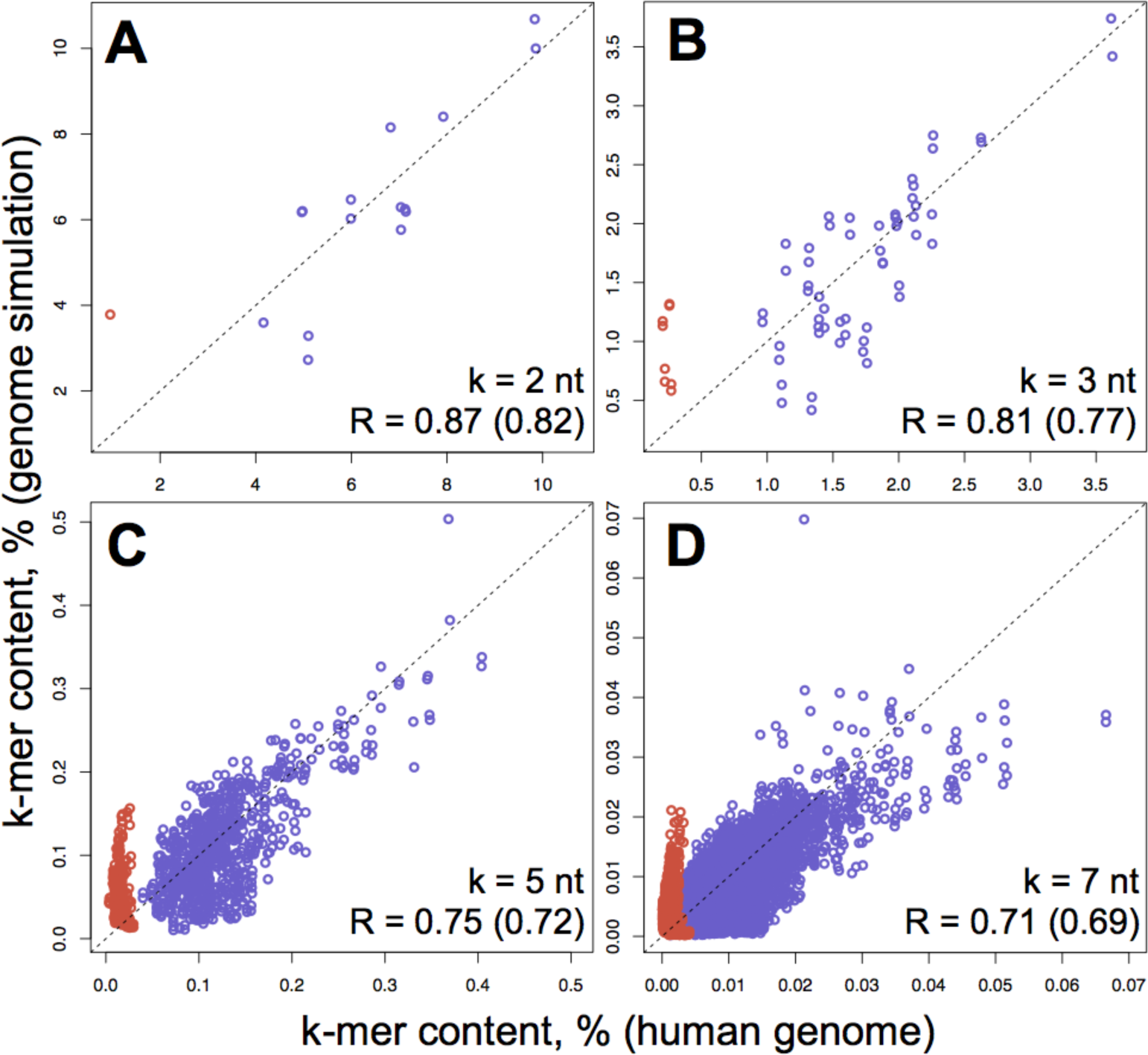
The comparison of the *in silico* (no strand-symmetries) evolved and real human genomes. The *in silico* genome is equilibrated here by using the raw Trek database but without accounting for the strand-symmetries of the 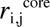 substitution rates (contrasting with **fig. 3D-G**). The plots **A-D** show the correlation of different k-mer contents in the equilibrated sequence with the corresponding content in the real human genome. The lengths of the k-mers along with the correlation coefficients are shown on the bottom right corners of the figure. Two correlation coefficients are shown with the exclusion and the inclusion (the value in the bracket) of the CpG containing oligomers (red points in the plots). The dashed lines depict the diagonals for the ideal match of the k-mer contents.

**Figure S9.**
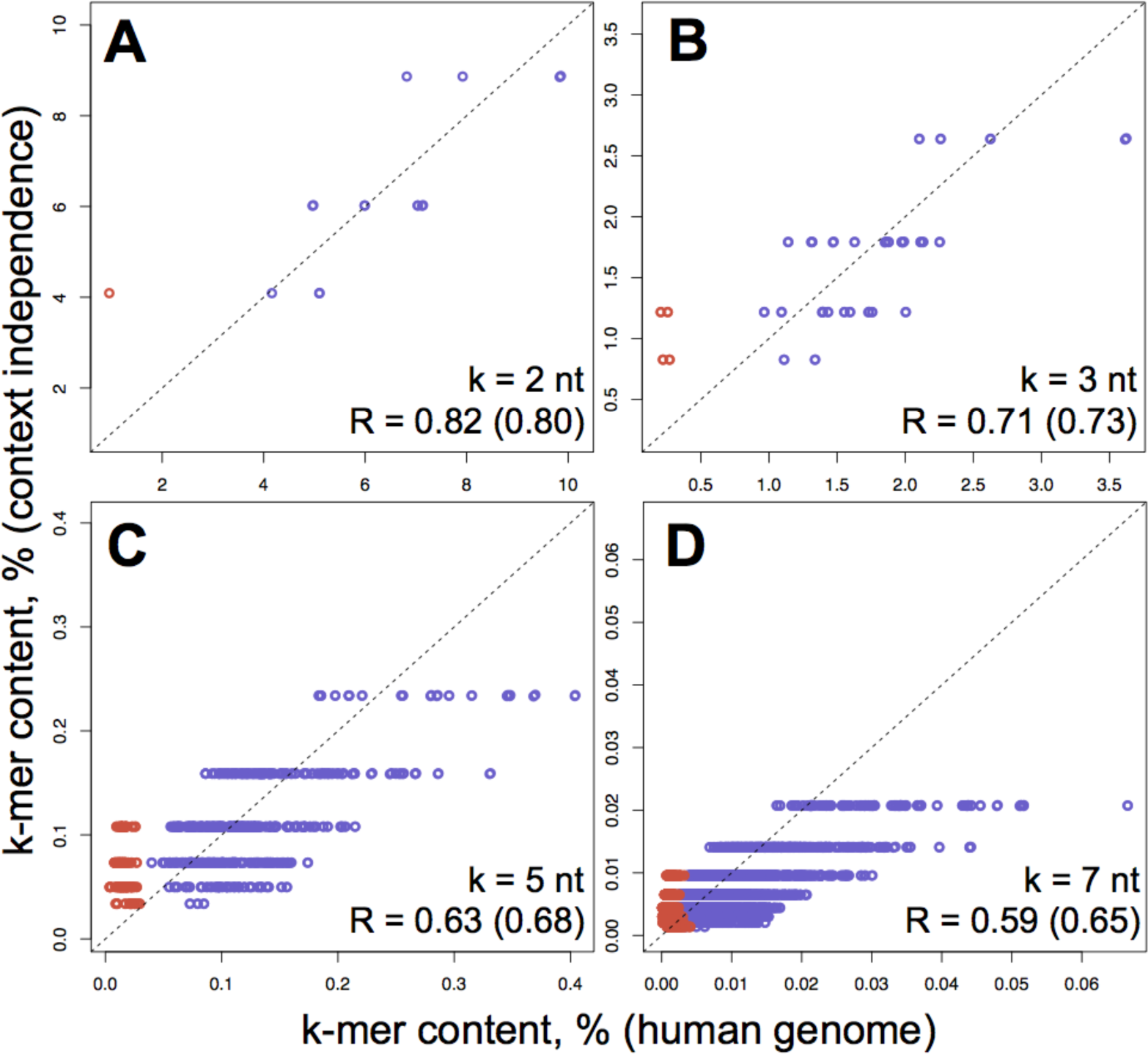
The oligomeric composition of a neighbour-invariant sequence. The oligomeric content of the human genome (x-axis) is compared to the content expected by a chance (y-axis) in a sequence that has the exact single-base composition as the human genome, but has substitution rates that are purely context independent. This corresponds to hypothetic genome simulation with perfectly correct, ideal 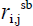 single-base rate constants, but without any 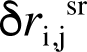 sequence-context dependence present. The lengths of the k-mers, along with the Pearson’s correlation coefficients without and with (the values in the brackets) the CpG containing oligomer data (red points) are shown on the bottom right corners of the plots. The correlation coefficients are notably smaller compared to the *in silico* sequence equilibrated based on the full set of context-dependent Trek 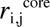 constants. The dashed lines depict the diagonals for the ideal match of the k-mer contents.

**Figure S10.**
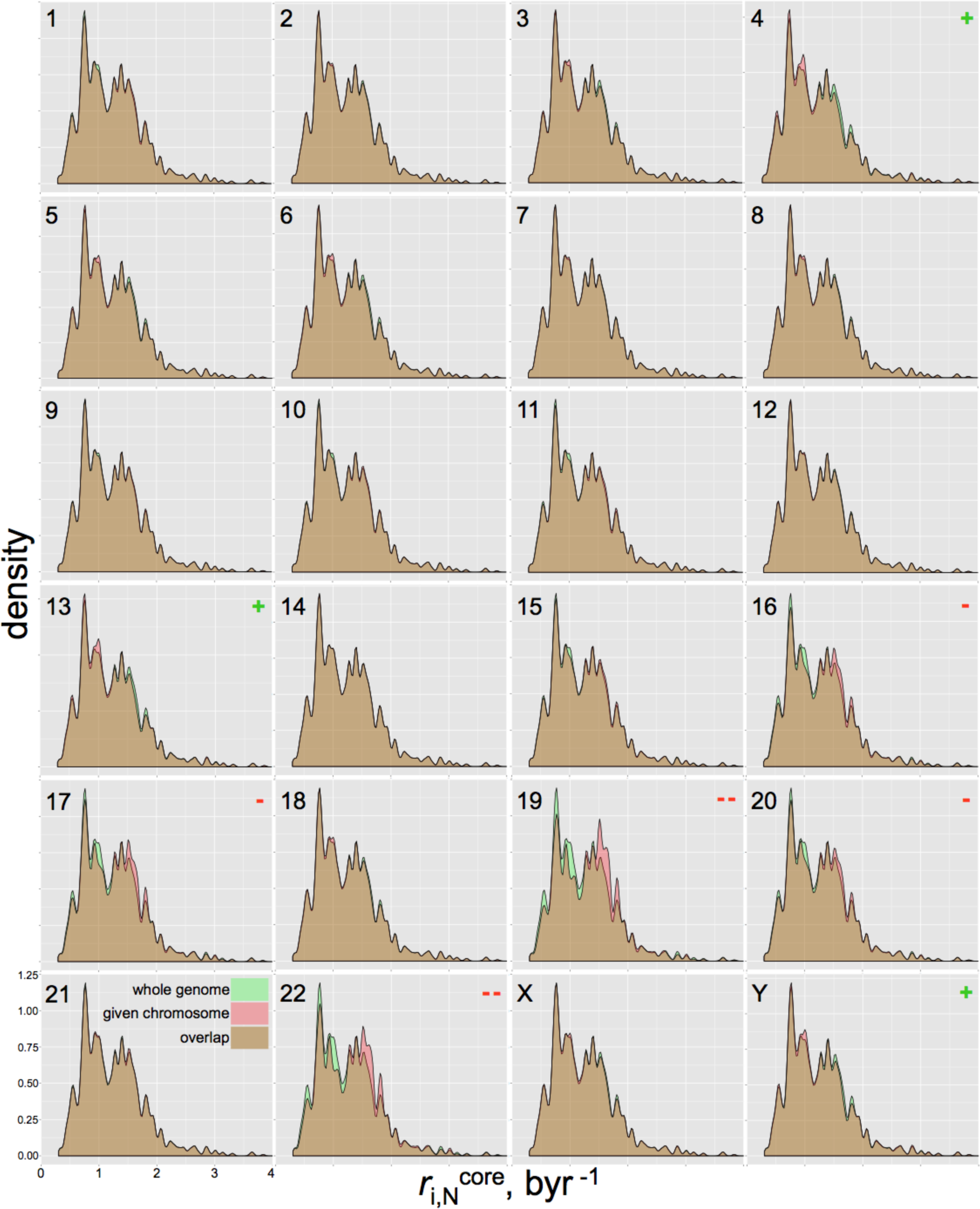
Basal substitution propensity (BSP) profiles of the human genome and its individual chromosomes. The plots show the density (kernel density estimate) distribution of the BSPs in the whole human genome (green), as compared to the individual chromosomes (red). The chromosome types are shown at the top-left corners of the plots. The overlaps of both distributions are in brown. The x-axis shows the BSP for the substitution to any other base 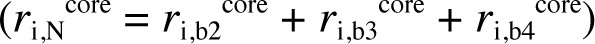, as inferred from mapping the positions with the context information (up to 7-mers) to the Trek database. Most of the chromosomes repeat the whole-genome substitution profile, with the significant exceptions noted for the chromosomes 19 and 22 that are relatively “destabilised”, in part due to their high G+C contents. The type (+ for stabilisation and – for destabilisation) of the difference between the profiles are shown at the top-right corners of the plots.

**Figure S11.**
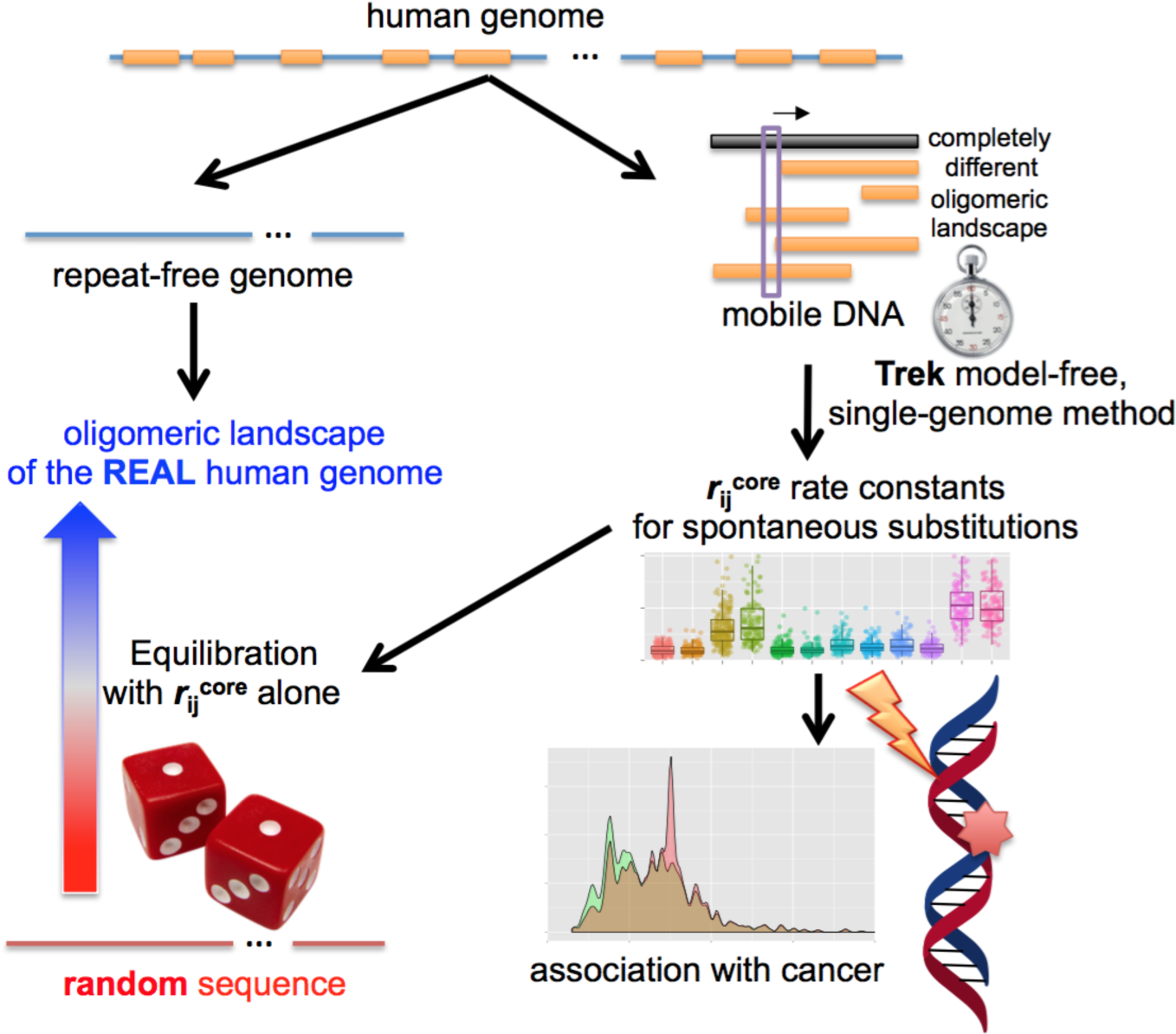
Schematic representation of the overall design and major outcomes of this work.

## DESCRIPTION OF THE SUPPLEMENTARY VIDEO AND DATA FILES

### supplementary_video_1.mov

#### Simulated evolution of a random genome guided solely by the 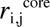 substitution rate constants

The starting 5mln-nt-long DNA sequence has 60% G+C content. The simulation goes on till the equilibration of the overall G+C content.

### supplementary_data.txt

#### data_1

Data on the Trek 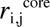 core substitution rate constants in the L1 reference sequence (L1Hs). The columns hold the substitution types, sequence positions, sequence contexts with 5 upstream and 5 downstream bases, 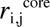 constants (byr^−1^), the correlation coefficient of the linear fit behind the 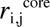 inference and the t-values.

#### data_2

Trek database processed with strand-symmetry consideration. The file contains all 7-mers, outlining the central base (i) where the rate constants (byr^−1^) are given for the core neutral substitutions into the 3 non-i bases (shown in an alphabetical order). The final columns hold the quality scores for 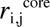 (actual sequence lengths matched with L1).

#### data_3

Trek database processed without strand-symmetry consideration. The file contains all 7-mers, outlining the central base (*i)* where the rate constants (byr^−1^) are given for the core neutral substitutions into the 3 non *-i* bases (shown in an alphabetical order). The final columns hold the quality scores for 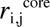 (actual sequence lengths matched with L1).

#### data_4

k-mer content for the repeat masked and unmasked versions of the human genome (RefSeq, hg19/GRCh37). The text file contains sections with headers showing the k-mer size and the genome masking status.

#### data_5

The full set of all 7-mer sequences with the respective cancer enrichment scores and basal substitution propensities. These data are based on the comparison of the COSMIC dataset with the 7-mer distribution in the human RefSeq. Only the non-SNP and non-coding sites undergoing single-nucleotide substitutions are taken from COSMIC (NCV set).

#### data_6

The GBM parameters that minimise the error of the test tree-based models, used to infer the optimal length window of neighbour effects. The columns represent the found best tuning parameters, along with the RMSE values for each *i→j* substitution type and window length.

## Acknowledgements

This research was supported by the Herchel Smith Fund and Cancer Research UK. SB is a Wellcome Trust Senior Investigator. We thank Dr. Chris Lowe for proofreading the manuscript.

